# How to describe and measure phenology? An investigation on the diversity of metrics using phenology of births in large herbivores

**DOI:** 10.1101/2021.05.17.444418

**Authors:** Lucie Thel, Simon Chamaillé-Jammes, Christophe Bonenfant

## Abstract

Proposed in 1849 by Charles Morren to depict periodical phenomena governed by seasons, the term “phenology” has spread in many fields of biology. With the wide adoption of the concept of phenology flourished a large number of metrics with different meaning and interpretation. Here, we first *a priori* classified 52 previously published metrics used to characterise the phenology of births in large herbivores according to four biological characteristics of interest: timing, synchrony, rhythmicity and regularity of births. We then applied each metric retrieved on simulation data, considering normal and non-normal distributions of births, and varying distributions of births in time. We then evaluated the ability of each metric to capture the variation of the four phenology characteristics *via* a sensitivity analysis. Finally, we scored each metric according to eight criteria we considered important to describe phenology correctly. The high correlation we found among the many metrics we retrieved suggests that such diversity of metrics is unnecessary. We further show that the best metrics are not the most commonly used, and that simpler is often better. Circular statistics with the mean vector orientation and mean vector length seems, respectively, particularly suitable to describe the timing and synchrony of births in a wide range of phenology patterns. Tests designed to compare statistical distributions, like Mood and Kolmogorov-Smirnov tests, allow a first and easy quantification of rhythmicity and regularity of birth phenology respectively. By identifying the most relevant metrics our study should facilitate comparative studies of phenology of births or of any other life-history event. For instance, comparative studies of the phenology of mating or migration dates are particularly important in the context of climate change.

## Introduction

In 1849, Charles Morren coined the term “phenology” to describe how periodical phenomena such as plant growth and reproduction are governed by the course of seasons (Morren 1849, see also Demarée 2011). With his observations he opened a new field of research and almost two centuries later the concept of phenology has become a cornerstone of ecology (Begon *et al*. 1986), used in plant and animal ecology simultaneously (Forrest and Miller-Rushing 2010). By describing when particular life-history events (*e*.*g*. flowering, parturition) occur in relation to the characteristics or states of the individual (*e*.*g*. size, age) as well as to environmental factors (*e*.*g*. photoperiod, predation risk) the concept of phenology is key to understanding the temporal cycles in the life history of species (Forrest and Miller-Rushing 2010). Nowadays, the term phenology is commonly employed to describe the temporal occurrence of many aspects of a species biology (*e*.*g*. moulting, migration, diapause in animals), but the phenology of reproduction (*e*.*g*. Sinclair *et al*. 2000, Rubenstein and Wikelski 2003, van den Hoff 2020) has attracted most interest. Reproductive phenology is an integral part of life history theory as it is at the heart of inter-generational trade-offs (*i*.*e*. between parents and offspring) and is a key factor of the reproductive success and fitness of the individuals (Stearns 1989, Forrest and Miller-Rushing 2010). On the one hand, the time of the year when most births occur is often linked to seasonal variations in food resources so that the flush of food resources matches the energetic needs of breeding, which ultimately improves the reproductive success of parents and the fitness of offspring (Plard *et al*. 2015). While on the other hand, the spread of birth dates in a year is supposed to reflect anti-predator strategies to reduce the mortality associated with predation (Darling 1938, Gosling 1969), but also many other social and biological mechanisms (Ims 1990), such as avoidance of male harassment undergone by females (Boness *et al*. 1995) or intra-specific competition between offspring (Hodge *et al*. 2011).

In most ecological studies, measurements and observations of phenology are frequently performed at the population level by characterising the temporal distribution of biological events (Visser *et al*. 2010). These rather complex and variable patterns are reduced to two main components: “timing”, the date at which the event of interest occurs, and “synchrony”, the spread of the dates at which the event occurs, *i*.*e*. the variability between individuals (Fig. 1). Stimulated by research on the effects of climate change on biodiversity (*e*.*g*. Crick and Sparks 1999, Parmesan 2007, Sarkar *et al*. 2019), the question of whether phenology is consistent or varies in time, both at individual and population levels, has received increased interest in recent years (*e*.*g*. Renaud *et al*. 2019). We therefore need to quantify two underappreciated properties of phenology: the consistency of the timing and synchrony (at the population scale) of the events from one reproductive season to the next. As these characteristics of phenology are not described by specific words yet, we suggest using “rhythmicity” and “regularity” to describe the consistency of timing and synchrony respectively (Fig. 1), in line with Newstrom’s terminology coined for tropical plants (Newstrom *et al*. 1994).

**Figure 1:**
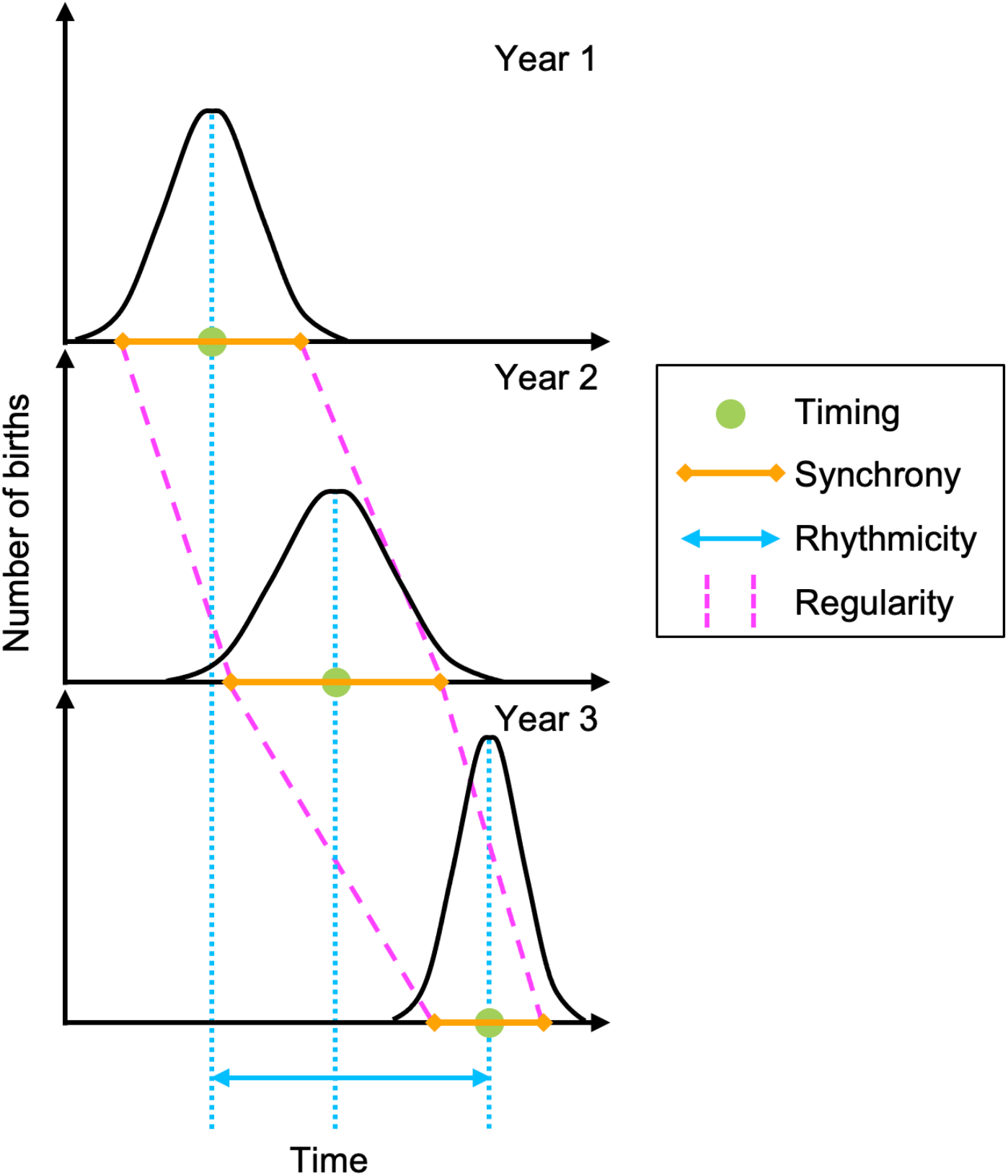
Four characteristics of phenology of births can be explored to fully describe phenology at the population scale: timing, synchrony, rhythmicity and regularity. Timing describes when within the year most births occur, synchrony illustrates whether females tend to give birth at the same time in a population in a given year, rhythmicity defines the consistency of timing between years, regularity refers to the consistency of synchrony between years. Green = timing, orange = synchrony, blue = rhythmicity, pink = regularity.

Despite appearing simple, the concept of phenology carries a lot of confusion in literature, both from a semantic and a descriptive point of view (Visser *et al*. 2010). Previous studies have explored phenology using a vast diversity of mathematical descriptors, many of which remain specific to a single study. This is problematic as well-defined, comparable and reliable descriptors of the temporal distribution of biological events are key to achieving meaningful comparisons of phenology patterns within or across species. English and colleagues reassessed the most influential factors of reproductive synchrony in large herbivores using the existing literature, but had to narrow their original data set because there was no standardised way of measuring and comparing synchrony across the studies (English *et al*. 2012). This large diversity of metrics is associated with a lack of widely accepted definitions or divergent definitions for the same word (see “seasonality” *sensu* Skinner *et al*. 2002 and Heideman and Utzurrum 2003), which further limits our ability to make meaningful comparisons (*e*.*g*. Ryan *et al*. 2007, Heldstab *et al*. 2018). As experimental studies are logistically challenging or virtually impossible to conduct with large species, the comparison of phenology patterns within a species living in contrasting environments or across species (Clauss *et al*. 2020) is of major importance to assess the role of explanatory factors accounting for the often marked variability in phenology reported in empirical studies (Rutberg 1987). Such comparative approaches (*sensu* Felsenstein 1985) indeed shed light on the ecological and evolutionary causes shaping the main stages of the life cycle of organisms (Bronson 1989).

Despite the increasing diversity of approaches to describe phenology, we found only a few attempts to compare phenology metrics and to provide advice on which one should be used preferentially according to the context of the study (Moussus *et al*. 2010, Landler *et al*. 2018). These initiatives are rare and we currently lack a comprehensive comparison of the metrics previously used to characterise phenology. The extent to which the different metrics capture the desired characteristics of the temporal distribution of events, or the sensitivity of those metrics to actual changes in phenology remain to be adequately assessed. Here, we propose such a comparison of metrics based on a literature survey of reproductive phenology in large herbivore species. We focus on the taxonomic group of the large herbivores as it has been studied in a number of species and at different locations (Rutberg 1987). As a result, we expect to find a wide variety of patterns of births and a wide diversity of metrics to describe them. We first clarify and formally define the four main terms describing phenology: timing, synchrony, rhythmicity and regularity, using our knowledge from the existing literature. We then conduct a comparative analysis of 52 metrics that have been used to quantify the different characteristics of phenology of births in large herbivores, highlighting their strengths and weaknesses. To conclude, we recommend one metric for each of the four main characteristics of phenology.

## Materials and methods

We conducted a quantitative comparison of a wide range of metrics used to analyse phenology in six steps. In Step 1, we recorded all metrics employed to measure phenology in a selection of papers that we considered representative of the study of phenology of births in large herbivores. In Step 2, we simulated contrasting phenology by varying independently the four parameters that determine timing, synchrony, rhythmicity and regularity of phenology of births (see details below). In Step 3, we calculated all metrics on the simulated phenology to understand how they compare and what characteristic of phenology they measure. In Step 4, we explored the similarities between metrics from a correlation matrix, and identified categories of metrics capturing the same characteristic of phenology. In Step 5, we evaluated the sensitivity of each metric to changes in the estimated parameter. In Step 6, we ranked each metric based on eight criteria that we considered important to identify robust and efficient metrics, but also meaningful from an ecological point of view (see Table 1 for a description of each criterion).

**Table 1:** Ordered list of the criteria used to evaluate the relevance of each metric describing phenology of births. Each criterion can be individually fully (score of 1) or partially (score of 0.5) validated or no (score of 0) by each metric. The value for the first four criteria (in bold type) should be > 0 to consider a metric to be possibly worthwhile and evaluate the remaining criteria. The sum of the value obtained for each criterion gives the relevance index of the metric (range between 0 and 8 points).

| Criterion | Description | Score |
| --- | --- | --- |
| **goodness** | Measures the parameter it is expected to measure | true = 1<br>false = 0 |
| **monotony** | Varies monotonically with the value of the parameter it is expected to measure, *i.e.* the sign of the slope coefficient is constant | true = 1<br>false = 0 |
| **saturation** | Does not saturate at the upper or lower boundary in a biological range of values (*e.g.* if a synchrony metric returned the same value when all births occurred during periods of various durations such as fifteen or thirty days, it was considered to saturate within a biologically realistic range of birth distributions because such distributions of births can be found in the wild) | true = 1<br>false = 0 |
| **strength**<sup>1</sup> | Is characterised by a strong relationship with the parameter it is expected to measure, *i.e.* is the scatter plot not too dispersed around the general trend of the relationship between the metric and the phenology characteristic, as an empirical approach of the predictive power? | high = 1<br>medium = 0.5<br>low = 0<br>(strength of the association) |
| normality | 1- Does not assume normally distributed birth dates; if false (assumes normality):<br>2- Is it robust to deviations to normality? *i.e.*, is the relationship between the metric and the parameter it is expected to measure conserved when births are not normally distributed | true = 1<br>false-true = 1<br>false-false = 0 |
| origin | Does not depend on the temporal origin set by the investigator | true = 1<br>false = 0 |
| linearity<sup>2</sup> | Is characterised by a linear relationship with the parameter it is expected to measure | type 1 = 1<br>type 2 and 3 = 0.5<br>type 4 = 0 |
| unicity | Gives a unique result | true = 1<br>false = 0 |

<sup>1</sup> high association: very small dispersion of points, medium association: small dispersion of points that does not prevent from detecting a trend, low association: dispersion of points large enough to prevent from detecting any trend, whatever the shape of the relationship (linear, but also sigmoid or quadratic for instance). <sup>2</sup> type 1 is a linear relationship, type 2 is a sigmoid-like relationship, type 3 is a quadratic-like relationship, type 4 is a binary relationship.

### Step 1: Retrieving and coding the different phenology metrics

We opportunistically searched the literature for articles focusing on the distribution of births in large herbivores using keywords such as “phenology”, “timing”, “synchrony”, “seasonality”, “period” or “season”, and using various sources such as search engines and the references in previously found articles. From these articles, published between 1966 and 2019, we recorded the metrics used to describe phenology of births at the population level. We stopped our search once the rate at which we discovered new metrics with additional papers became negligible.

We *a priori* classified each metric into one out of four categories based on our understanding of the original description and formula of the metric (Fig. 1): (1) *timing* metrics, defining when within the year most births occur, (2) *synchrony* metrics, defining whether females tend to give birth at the same time in a population in a given year, (3) *rhythmicity* metrics, defining the consistency of timing between years, (4) *regularity* metrics, defining the consistency of synchrony between years. In the literature, the term “seasonality” can be used to describe the location of births in the year (*i*.*e*. timing, *e*.*g*. in Sinclair *et al*. 2000), the duration of birth period (*i*.*e*. synchrony, *e*.*g*. in Zerbe *et al*. 2012), and even the fact that births occur at the same period of the year every year (*i*.*e*. rhythmicity and/or regularity, *e*.*g*. in Heideman and Utzurrum 2003). However, this term is initially used to describe the cyclical nature of the environment in a wider range than the study of birth phenology (Visser *et al*. 2010). Thus, it should be used to describe organisms’ phenology only when a direct relationship between periodic environmental phenomena and the cycle of the organism at stake has been demonstrated, which is not always the case in phenology studies. For this reason, we suggest using the term “seasonality” only to describe the cyclicity of the environment and prefer the use of neutral terms such as those we introduced in this paper to describe phenology of births: rhythmicity and regularity.

Forty-seven articles (Supporting information 1) presented at least one mathematically-defined phenology metric yielding 52 different metrics. In order to compare metrics quantitatively, we slightly tweaked some of them: when the metric was a boolean (true/false) variable based on the significance of a statistical test (*n* = 9 metrics), we used the value of the test statistic as output metric, thereby allowing us to investigate how the statistic was influenced by the value of phenology parameters (see details in Supporting information 2). All metrics could be coded in R software (R Core Development Team 2019) except one, for which Perl was used (www.perl.org).

### Step 2: Simulating phenology of births

We simulated phenology of births from statistical distributions with known parameters (Supporting information 2) to assess what characteristic of phenology (timing, synchrony, rhythmicity, regularity) each metric would capture, their sensitivity to changes into these four key characteristics of interest, and the correlation between the 52 metrics. We simulated the distributions of births over a year as most large herbivores breed once a year. This choice does not limit the generality of our results: for species breeding more than once per year (*e*.*g*. small species with short gestation length such as dikdik *Rynchotragus (Madoqua) kirki*, Sinclair *et al*. 2000), the same metrics may be applied on sub-periods of time, each displaying only one birth peak (see Heideman and Utzurrum 2003 for a similar approach in bats).

Each simulated phenology was generated by randomly distributing births in time, following a normal distribution. We distributed *n =* 1000 births within a year of 365 days, repeated over 10 years (see why in “Material and Methods” section, step 3). We changed four parameters independently to modify the distribution of births: the mean day of birth for a given year (*mean*), the standard deviation of the distribution of births for a given year (*sd*), the range over which the mean birth date can vary across years (*Δmean*), and the range over which the standard deviation can vary across years (*Δsd*). Each parameter varied in a range from a minimum to a maximum value and was incremented with a constant step (Supporting information 2). Choosing the value of these parameters allowed us to simulate changes in the timing, synchrony, rhythmicity and regularity of the phenology of births independently. As the simulated phenology of births relied on random draws, the actual values of parameters in the simulated distribution of births could differ from the theoretical values used in the simulation algorithm. We used the realised values of the distribution parameters in the following analyses. Note that we replicated the same analyses using non-normal distributions of births (*i*.*e*. skewed normal, bimodal, Cauchy, and random distributions) to cover the variety of empirical distributions of births observed in *natura* and assess robustness to non-normality (Supporting information 4). We performed all simulations using the R software and made the code available on GitHub (https://github.com/LucieThel/phenology-metrics).

### Step 3: Computing the phenology metrics from simulated patterns of births

Among the 52 phenology metrics we analysed, most applied to a single year, but others required two or more years of data to be computed (see the complete list in Supporting information 3). As we aimed to compute all 52 metrics, we chose to simulate annual distributions of births over 10 consecutive years by default. For each simulation, we used data from the first year to compute metrics requiring only one year of data (*n* = 33 metrics), data from the first two years for metrics requiring two years of data (*n* = 9 metrics), and data from the whole simulation for the other metrics (*n* = 10 metrics).

### Step 4: Comparing the metrics

With the results from Step 3, we computed the global correlation matrix between all pairs of metrics using Pearson correlations. We then identified groups of strongly correlated metrics from the pairwise correlation coefficients and assigned each metric to one or several of the four characteristics of phenology it was best related to. We compared this categorisation with our *a priori* classification of the metrics. This step enabled us to check our intuitive classification of the metrics in addition to revealing whether some metrics could incidentally capture several aspects of the distribution of births at once.

### Step 5: Estimating the sensitivity of the metrics

For each metric, we performed a sensitivity analysis by quantifying the observed variation of each metric with a fixed variation in the characteristic of phenology it was previously associated with in Step 4. We did this by computing, for each possible pair of simulations within the set of all simulations performed, the proportional difference between the realised values of the phenology parameter of interest of the two simulations, and the proportional difference between the values of the metric of interest of the same two simulations. In each case the proportional difference was calculated as *[(Value*_*max*_ *– Value*_*min*_*) / Value*_*min*_*] * 100*. This formulation allowed us to work with positive values only as we were interested in the amplitude but not in the direction of the differences.

### Step 6: Scoring metrics

Finally, as there were too many different metrics, we were unable to discuss the pros and cons for each of them. We chose instead to provide guidance about the usefulness of the different metrics by scoring them according to a set of eight criteria that we considered as important behaviour for a metric to be relevant (Table 1). Having systematic criteria helped us to minimise the subjectivity of the scoring so we ranked the metrics from 0 (not advised) to 8 (strongly advised) according to the number of criteria they fulfilled. The proposed criteria (Table 1) consisted in verifying if 1) the metric varied according to the phenology characteristic it was supposed to measure, 2) the variation of the metric according to the phenology characteristic was monotonous, 3) the relationship with the characteristic of phenology was strong (visual assessment of the association between the computed statistic and the phenology characteristic), 4) the metric did not saturate within a biologically realistic range of distributions of births. We considered that metrics with scores < 4 for which the first four essential criteria were not validated should not be advised. If those four criteria were satisfied, we evaluated an additional set of four criteria (normality, independence of the temporal origin, linearity and unicity of the output, see Table 1 for a detailed description). All criteria were scored from visual inspection of the results by one of us (LT).

## Results

The mean number of metrics used in each paper was 3.8 ± 2.1 sd (*range* = 1 - 8). Eleven metrics were *a priori* associated with timing, 25 with synchrony, 10 with rhythmicity and five with regularity. We did not classify one metric because it could either be a rhythmicity or regularity metric *a priori*. Those metrics were based on descriptive statistics, circular statistics, statistical tests or statistical modelling such as general linear models. The unit of the metrics were date, duration, counts (*e*.*g*. a number of births), binary classification (*i*.*e*. if a given condition was satisfied or not), or unitless indices (Supporting information 3).

The correlation matrix (Step 4) revealed groups of metrics that were highly correlated and thus reflected the same characteristic of phenology (Fig. 2). Five groups were clearly identifiable, representing timing metrics (Fig. 2 - box 1), synchrony metrics (Fig. 2 - boxes 2 and 5), rhythmicity metrics (Fig. 2 - box 3), and regularity metrics (Fig. 2 - box 4). The two groups of metrics measuring synchrony had highly but negatively correlated values (Fig. 2 - box 6). This indicated that all metrics of the two groups captured synchrony correctly, however, in an opposing way. Three metrics were singular and were associated with neither of the five groups. The metric which compares the slope coefficients of linear models describing the log percent of cumulative births (“*splcomp*”) should measure regularity, but it rather correlated better with synchrony metrics. The metric which evaluates the duration between the first birth dates of two reproductive cycles (“*diffbgper*”), an assessment of rhythmicity, correlated well with both rhythmicity and regularity metrics. Seven other metrics had a detectable relationship with at least one of the three remaining phenology characteristics in addition to the relationship with the phenology characteristic they were supposed to quantify (Supporting information 3 and 5).

**Figure 2:**
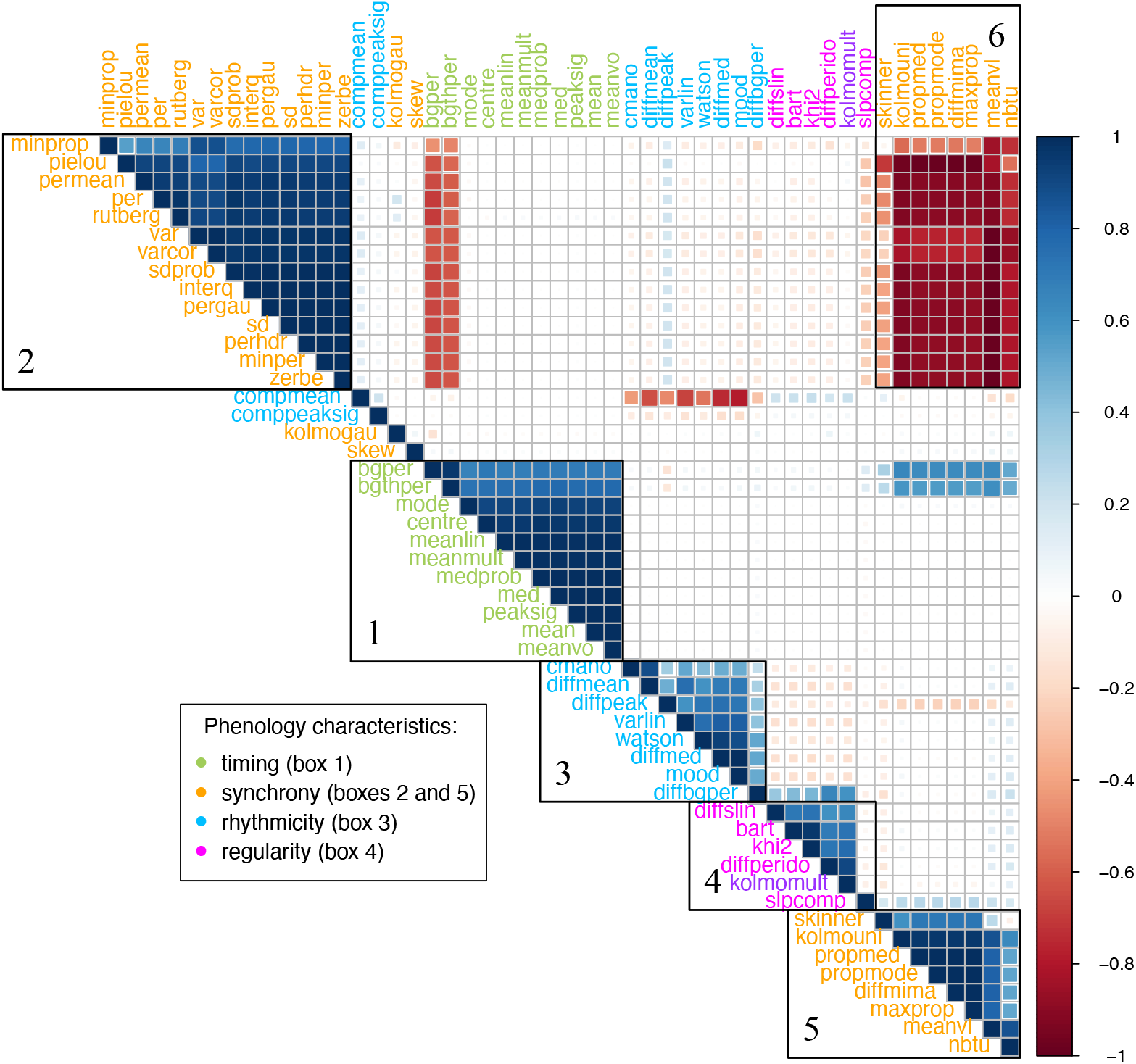
Correlation matrix between all pairs of metrics, using Pearson correlations (n = 51, “rayleigh” removed because because of no observed variation). It was not possible to classify “kolmomult” a priori in rhythmicity or regularity metrics, as it compares the complete distribution of births between two years. Box 6 highlights the high but negative correlation between the two groups of metrics measuring synchrony (boxes 2 and 5). Green = timing metrics, orange = synchrony metrics, blue = rhythmicity metrics, pink = regularity metrics. Note the high negative correlation between “compmean” and the other rhythmicity metrics, highlighting that it is also a rhythmicity metric.

The sensitivity of the metrics to the simulated variation of the phenology characteristics (Step 5) differed markedly between metrics, especially in synchrony and regularity metrics (Fig. 3 and Supporting information 5). The proportion of variation of the metrics for a 10 % variation of the associated parameter ranged from 14 % to 33 % for timing metrics, from 0 % to 139 % for synchrony metrics, from 0 % to 471 % for rhythmicity metrics and from 0 % to 138 % for regularity metrics. The variation of almost all timing, rhythmicity and regularity metrics according to variations of their associated parameter was highly homogeneous. Synchrony metrics were less homogeneous, certainly due to the fact that those metrics were the most numerous and based on more diverse methods (proportion of variation, integrative indexes or moments of the distribution of births, for instance). The metrics that were singular in the correlation matrix were clearly visible in the heat maps, characterised by erratic or non-existent variations (*e*.*g*. skewness of the birth distribution “*skew*”, and comparison of mean date of births “*compmean*”).

**Figure 3:**
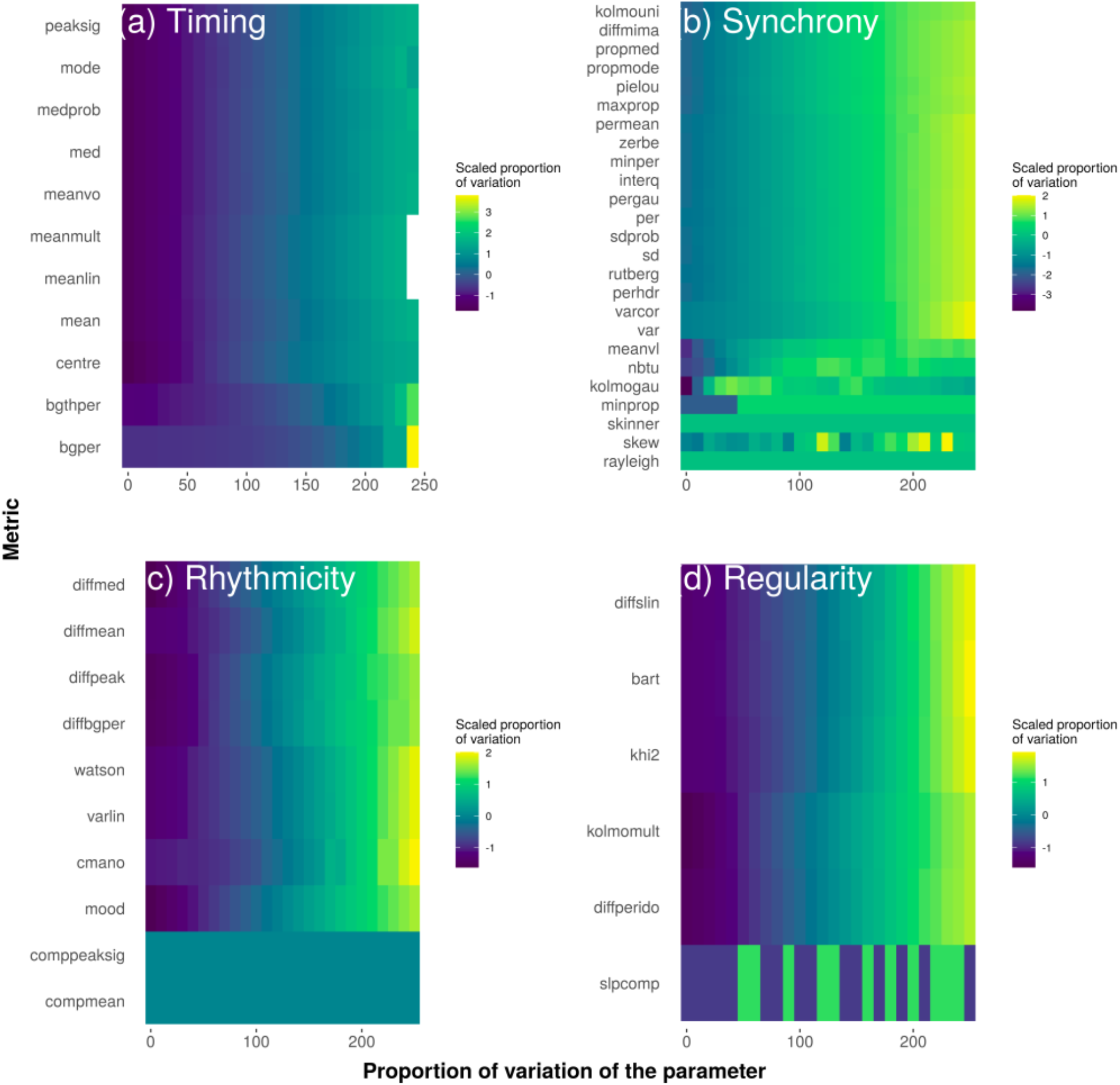
Heat maps representing the (scaled) proportion of variation of the metric in relation to the proportion of variation of the parameter of phenology (sensitivity analysis): a) timing metrics according to the mean birth date for a given year (mean, n = 11), b) synchrony metrics according to the standard deviation of the distribution of births for a given year (sd, n = 25), c) rhythmicity metrics according to the range over which the mean birth date can vary across years (Δmean, n = 10), d) regularity metrics according to the range over which the standard deviation of the distribution of births can vary across years (Δsd, n = 6).

*Colours in the heat maps reflect the proportion of variation of each metric according to the proportion of variation of the phenology parameter, normalised for each metric using all values of the metric obtained across all simulations. We normalised the sensitivity of each metric individually to prevent the representation of the large variation of some metrics to hide the smaller but meaningful variations of other metrics to be visible. Metrics characterised by a large colour gradient vary widely in response to the variation of the parameter of phenology they measure. Metrics with a smoothed colour transition vary regularly in response to the variation of the parameter of phenology they measure. To the contrary, metrics characterised by sudden and/or random colour transitions vary inconsistently in response to the variation of the parameter of phenology we changed*.

The same analyses conducted on the basis of non-normal distributions led to similar observations in the case of asymmetric distributions (skewed normal, bimodal and Cauchy distributions). The correlation matrices showed similar patterns of correlations between the metrics, and the metrics varied analogously according to the variation of the *mean, sd, Δmean* and *Δsd* of the distributions for normal and asymmetric distributions either (see Supporting information 4 for a detailed analysis). Nevertheless, it is worth noting that a very limited number of metrics depending on the skewness of the distribution did not perform as well with the normal distribution than with asymmetric distributions. On the contrary, metrics depending on the presence of a period without any birth did not perform as well with non-normal distributions than with a normal distribution. In the case of a random distribution, no clear correlations between metrics nor relationships between the metrics and the four parameters of the distribution were detectable, except for some rare synchrony and timing metrics (Supporting information 4).

The relevance score of the metrics (step 6) varied between 0 and 8, covering the complete range of variation possible (Fig. 4) and we list, for each phenology characteristic, the metrics we identified as “best” (Table 2). Our classification also revealed what could be considered as ineffective (*score* = 0, *n* = 4) and poor metrics (*score* ∈ [0; 4[, *n* = 14). All the timing metrics reached excellent scores above 6. Nevertheless, the mean vector orientation (“*meanvo*”) was the best metric, fulfilling all our criteria with a score of 8 (Fig. 4). Three metrics provided a very good assessment of the synchrony of births with a score of 7.5: the evenness index (“*pielou*”), the mean vector length (“*meanvl*”) and the comparison of the distribution of births to a uniform distribution (“*kolmouni*”) (Fig. 4). The best metric to quantify rhythmicity measured the time elapsed between the median birth dates of two years (“*diffmed*”), with a score of 7 (Fig. 4). It is worth noting that the non-parametric Mood test (“*mood*”) provides a statistical assessment of whether “*diffmed*” differs from 0. The non-parametric Mood test (“*mood*”) obtained a marginally lower score (6.5, Fig. 4) than “*diffmed*” only because of a slight non-linearity in the relationship between simulation parameter values and the metric’s statistics. Altogether, we therefore considered that “*mood*” could be very useful to measure rhythmicity. One metric quantifying regularity stood out from the others according to our criteria: the non-parametric Kolmogorov-Smirnov test (“*kolmomult*”), which compares two birth distributions (*score* = 7.5, Fig. 4).

**Table 2:** List of the metrics considered as the best metric, for each characteristic of the phenology of births (timing, synchrony, rhythmicity, regularity).

| Phenology characteristic | Metric | Complete name | Description | Reference |
| --- | --- | --- | --- | --- |
| Timing | *meanvo* | mean vector orientation | evaluates mean vector orientation of the birth distribution | Paré *et al.* 1996 |
| Synchrony | *meanvl* | mean vector length | evaluates mean vector length of the birth distribution | Paré *et al.* 1996 |
| Rhythmicity | *mood* | Mood test | compares median birth dates between two years | Berger and Cain 1999 |
| Regularity | *kolmomult* | Two-sample Kolmogorov-Smirnov test | compares birth distributions between two years | Green and Rothstein 1993 |

**Figure 4:**
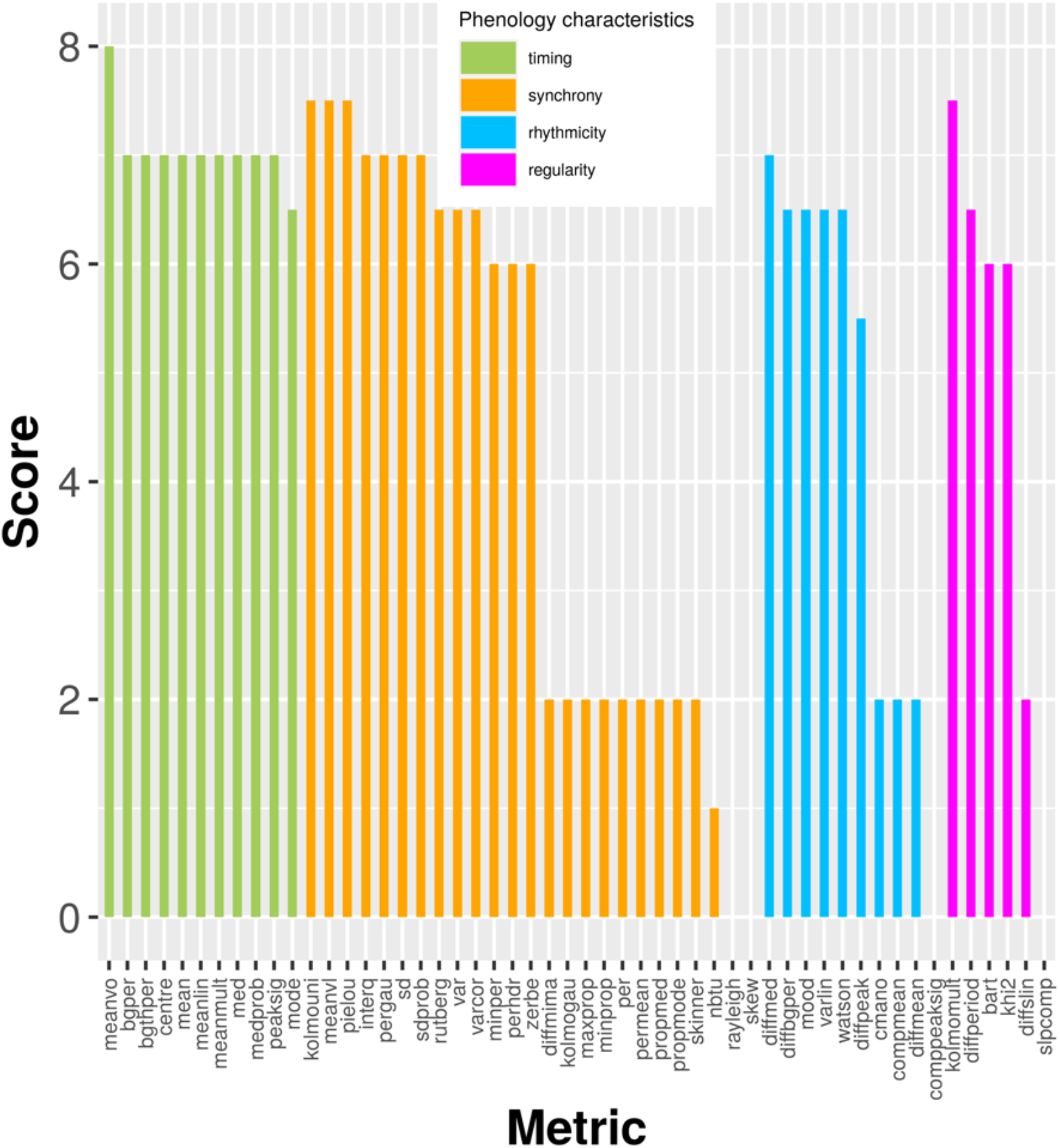
Score obtained by each phenology metric (n = 52) according to the eight criteria used to assess its relevance to characterise the four main characteristics of birth phenology (goodness, monotony, saturation, strength, normality, origin, linearity and unicity, as defined in Table 1). Green = timing metrics, orange = synchrony metrics, blue = rhythmicity metrics, pink = regularity metrics.

## Discussion

With more than fifty metrics used to describe and analyse the distribution of births in large herbivores since 1966, our survey of the literature clearly illustrates the diversity of approaches, even when focusing on a specific taxonomic group. Although the choice of a metric is most of the time justified, either to answer a specific ecological question or on statistical grounds, the lack of consensual methods to quantify phenology makes comparisons across species or populations difficult at best, if possible at all. Our simulation study suggests that such a diversity of metrics may cause confusion and be unnecessary as we were able to identify a reduced set of simple metrics that works well to measure the different characteristics of phenology. Moreover, we believe our work can also provide insights into how to analyse phenology of other traits than birth dates, such as migration dates of birds or flowering dates (Moussus *et al*. 2010).

Many of the metrics we retrieved can be organised into four main categories, each one capturing a particular characteristic of phenology: timing, synchrony, rhythmicity and regularity. Of course, metrics belonging to the same category are not perfectly equivalent and interchangeable (Fig. 2, see also a discussion comparing “*zerbe*” and “*rutberg*” metrics in Zerbe *et al*. 2012). For instance, the correlations between timing metrics range between 0.68 and 1.00. The difference among metrics is more pronounced in the synchrony category with correlations ranging from 0.05 to 1.00 (excluding “*kolmogau*” and “*skew*” metrics that appear as singularities in the correlation matrix, Fig. 2). How different characteristics of phenology are measured can lead to dependency between one another and this could explain the confusions found in the literature between timing and synchrony through terms such as “period” or “season” of births. Indeed, several of the metrics we tested vary not only according to the phenology characteristic they were used to measure, but also according to other characteristics of the phenology (*n* = 8 metrics). For instance, we show a strong correlation between metrics that evaluate the start of the birth period (*i*.*e*. timing metrics “*bgper*” and “*bgthper*”) and the synchrony metrics in general. This association between different types of metrics arises when the standard deviation of the simulated distributions of births increases (while the mean is fixed), leading to earlier births (Fig. 2).

We attempted to identify what metrics could be the most suitable for measuring timing, synchrony, regularity and rhythmicity of phenology by scoring them according to what we subjectively considered as the main suitable properties. We considered that a good metric should not be restricted to one kind of pattern (*e*.*g*. unimodal) as the distribution of births is not necessarily known a priori and may change between years due to ecological factors (see Adams and Dale 1998 for instance). Slightly more than 10 % of the metrics theoretically require normally distributed dates of birth to work well (based on the metrics for which this criterion was evaluated, Supporting information 3). We showed these metrics are generally robust to deviations from normality so this assumption does not limit their application to most data. The metrics should also be independent of the temporal origin set by the investigator, as the favourable periods for reproduction cycle differ between species and populations (*e*.*g*. mountain sheep *Ovis spp*. inhabiting desert and alpine ecosystems, Bunnell 1982). Using the calendar year would be biologically meaningless and will create artificial patterns of births by splitting the distribution around the end of the year. We identified six metrics independent of the temporal origin: the day with the highest number of births (“*mode*”), the evenness index (“*pielou*”), the mean vector orientation and length form the circular statistics (“*meanvl*” and “*meanvo*” respectively), and the non-parametric Kolmogorov-Smornov test comparing a birth distribution to a uniform distribution or another birth distribution (“*kolmouni*” and “*kolmomult*” respectively). Circular statistics could be favoured to answer the difficulties linked to the selection of temporal origin, as it is frequently done in primate literature (*e*.*g*. Di Bitetti and Janson 2000). Notwithstanding such limitations, we found several metrics that met our expectations of a good metric for each phenology characteristic (Table 2 and Figure 4).

On the other side a few metrics should not be recommended to describe phenology of births. The evaluation of rhythmicity describing the evolution of the mean dates of births of several years with a linear regression (“*diffmean*”), or the quantification of synchrony through the duration of the period gathering at least a certain percent of births (“*nbtu*”) are not to be advised. In addition to undesirable statistical properties, these metrics fail to capture the changes in the phenology parameter adequately. The metric “*nbtu*” varied non-monotonously with the level of synchrony of the birth phenology. Similarly, the duration between first and last birth to measure synchrony (“*per*”) plateaued for a range of biologically realistic values, what limits its usability in a wide range of ecological conditions.

Overall, some phenology characteristics have been more consistently evaluated across studies, a fact illustrated by the number of metrics of each category used in more than two papers (*n* = 5, 7, 2 and 0 for timing, synchrony, rhythmicity and regularity respectively, see Supporting information 6). If timing and synchrony of births are the easiest and most frequent characteristics of phenology estimated and compared, only a handful of metrics evaluates rhythmicity and regularity of the phenology of births across the years. Sound analysis of rhythmicity and regularity indeed requires many years of data which may not be available as such data is costly and time-consuming to collect (Kharouba and Wolkovich 2020). Moreover, scientists are less interested in timing and synchrony consistency *per se* than in the relationship between timing and synchrony, and ecological or environmental factors such as temperature, rainfall or spring snow cover (Paoli *et al*. 2018). Our study shows that the rhythmicity and regularity metrics currently available are only moderately correlated, particularly when they are used to describe birth distributions that are not normally distributed (Supporting information 4). Capturing the temporal variation of phenology across years appears difficult and requires thoughtful selection and interpretation of the used metric. Standardised and relevant statistical tools are needed to quantify regularity and rhythmicity of phenology, and to test their hypothetical responses to global changes. This study should help in this.

Although we show that the assumption of a normal distribution or another bell-shaped (asymmetric or not) distribution mimicking those found in *natura* (*e*.*g*. skewed normal, bimodal or Cauchy distribution) has no major consequences on our conclusions (Supporting information 4), this is not true when there is no clear pattern in the distribution of births. Indeed, most metrics give inconsistent and unreliable results when applied to birth dates randomly distributed within the year (Supporting information 4), a pattern that has been documented in some populations of large herbivores living in the southern hemisphere (Sinclair *et al*. 2000). Describing random patterns using the metrics presented here is unlikely to be useful because biologically meaningless: when births occur year-round, the timing and rhythmicity are meaningless as they cannot reduce to one or two summarising statistics. Using evenness indexes such as “*pielou*” could at least provide a quantification of the heterogeneity of the distribution of births.

In conclusion, we recommend using the circular mean vector orientation (“*meanvo*”) to describe timing and the circular mean vector length (“*meanvl*”) to describe synchrony, because both are not influenced by the temporal origin set by the investigator. We recommend using the underused Mood test which statistically compares the median birth dates (“*mood*”) to describe rhythmicity and the Kolmogorov-Smirnov test which statistically assesses if two birth distributions are similar to describe regularity (“*kolmomult*”, see Table 2 and Supporting information 3 for a formal description of those metrics). Being non-parametric tests, they are applicable in a wide range of distributions as frequently observed in large herbivore populations.

## Acknowledgments

The authors thank Simon Penel for his help with Perl Programming Language, and Marcus Clauss for providing the code of one of the metrics. The authors thank two anonymous reviewers and the subject editor for their comments on a previous version of the manuscript, and Marion Valeix for her comments on early drafts of the manuscript. They deeply appreciated the insightful comments, editorial suggestions and proofreading by Anna Cryer, Leif Egil Loe and Primaëlle Fusto. This work was performed using the computing facilities of the CC LBBE/PRABI and the IFB Cloud.

## Declarations

### Funding

This work was supported by a grant from the “Ministère Français de l’Enseignement Supérieur, de la Recherche et de l’Innovation” through the “Ecole Doctorale E2M2” of the “Université Claude Bernard Lyon 1”.

### Authors’ contributions

LT, CB and SCJ conceived the ideas and designed the methodology; LT performed data collection and analysed the data; LT wrote the first draft of the manuscript, which was then edited by all authors. All authors gave final approval for publication.

### Data availability statement

Code available on GitHub: https://github.com/LucieThel/phenology-metrics.

## Notes

https://github.com/LucieThel/phenology-metrics

### Competing Interest Statement

The authors have declared no competing interest.

## References

Adams, L. G. and Dale, B. W. 1998. Timing and synchrony of parturition in Alaskan caribou. - J. Mammal. 79: 287–294.

Begon, M. et al. 1986. Ecology. Individuals, populations and communities. - Blackwell scientific publications.

Berger, J. and Cain, S. L. 1999. Reproductive synchrony in brucellosis-exposed bison in the southern Greater Yellowstone Ecosystem and in noninfected populations. - Conserv. Biol. 13: 357–366.

Boness, D. J. et al. 1995. Does male harassment of females contribute to reproductive synchrony in the grey seal by affecting maternal performance? - Behav. Ecol. Sociobiol. 36: 1–10.

Bronson, F. H. 1989. Mammalian reproductive biology. - University of Chicago Press.

Bunnell, F. L. 1982. The lambing period of mountain sheep: synthesis, hypotheses, and tests. - Can. J. Zool. 60: 1–14.

Clauss, M. et al. 2020. Basic considerations on seasonal breeding in mammals including their testing by comparing natural habitats and zoos. - Mamm. Biol.: 1–14.

Crick, H. Q. P. and Sparks, T. H. 1999. Climate change related to egg-laying trends. - Nature 399: 423.

Darling, F. 1938. Bird flocks and the breeding cycle. - Cambridge University Press.

Demarée, G. R. 2011. From “Periodical Observations” to “Anthochronology” and “Phenology”- the scientific debate between Adolphe Quetelet and Charles Morren on the origin of the word “Phenology.” - Int. J. Biometeorol. 55: 753–761.

Di Bitetti, M. S. and Janson, C. H. 2000. When will the stork arrive? Patterns of birth seasonality in neotropical primates. - Am. J. Primatol. Off. J. Am. Soc. Primatol. 50: 109–130.

English, A. K. et al. 2012. Reassessing the determinants of breeding synchrony in ungulates. - PLoS One 7: e41444.

Felsenstein, J. 1985. Phylogenies and the comparative method. - Am. Nat. 125: 1–15.

Festa-Bianchet, M. 1988. Birthdate and survival in bighorn lambs (Ovis canadensis). - J. Zool. 214: 653–661.

Forrest, J. and Miller-Rushing, A. J. 2010. Toward a synthetic understanding of the role of phenology in ecology and evolution. - Philos. Trans. R. Soc. B Biol. Sci. 365: 3101–3112.

Gosling, L. M. 1969. Parturition and related behaviour in Coke’s hartebeest, Alcelaphus buselaphus cokei Günther. - J. Reprod. Fertil. Suppl. 6: 265–286.

Green, W. C. H. and Rothstein, A. 1993. Asynchronous parturition in bison: implications for the hider-follower dichotomy. - J. Mammal. 74: 920–925.

Heideman, P. D. and Utzurrum, R. C. B. 2003. Seasonality and synchrony of reproduction in three species of nectarivorous Philippines bats. - BMC Ecol. 3: 11.

Heldstab, S. A. et al. 2018. Geographical Origin, Delayed Implantation, and Induced Ovulation Explain Reproductive Seasonality in the Carnivora. - J. Biol. Rhythms 33: 402–419.

Hodge, S. et al. 2011. Reproductive competition and the evolution of extreme birth synchrony in a cooperative mammal. - Biol. Lett. 7: 54–56.

Ims, R. A. 1990. The ecology and evolution of reproductive synchrony. - Trends Ecol. Evol. 5: 135–140.

Kharouba, H. M. and Wolkovich, E. M. 2020. Disconnects between ecological theory and data in phenological mismatch research. - Nat. Clim. Chang.: 1–10.

Landler, L. et al. 2018. Circular data in biology: advice for effectively implementing statistical procedures. - Behav. Ecol. Sociobiol. 72: 128.

Morren, C. 1849. Le globe, le temps et la vie. - Bulletins de l’Académie royale des Sciences, des Lettres et des Beaux-Arts de Belgique 16: 660–684.

Moussus, J.-P. et al. 2010. Featuring 10 phenological estimators using simulated data. - Methods Ecol. Evol. 1: 140–150.

Newstrom, L. E. et al. 1994. A new classification for plant phenology based on flowering patterns in lowland tropical rain forest trees at La Selva, Costa Rica. - Biotropica: 141–159.

Paoli, A. et al. 2018. Winter and spring climatic conditions influence timing and synchrony of calving in reindeer. - PLoS One 13: e0195603.

Paré, P. et al. 1996. Seasonal reproduction of captive Himalayan tahrs (Hemitragus jemlahicus) in relation to latitude. - J. Mammal. 77: 826–832.

Parmesan, C. 2007. Influences of species, latitudes and methodologies on estimates of phenological response to global warming. - Glob. Chang. Biol. 13: 1860–1872.

Plard, F. et al. 2015. The influence of birth date via body mass on individual fitness in a long-lived mammal. - Ecology 96: 1516–1528.

Renaud, L.-A. et al. 2019. Phenotypic plasticity in bighorn sheep reproductive phenology: from individual to population. - Behav. Ecol. Sociobiol. 73: 50.

Rubenstein, D. R. and Wikelski, M. 2003. Seasonal changes in food quality: a proximate cue for reproductive timing in marine iguanas. - Ecology 84: 3013–3023.

Rutberg, A. T. 1987. Adaptive hypotheses of birth synchrony in ruminants: an interspecific test. - Am. Nat. 130: 692–710.

Ryan, S. J. et al. 2007. Ecological cues, gestation length, and birth timing in African buffalo (Syncerus caffer). - Behav. Ecol. 18: 635–644.

Sarkar, U. K. et al. 2019. Climato-environmental influence on breeding phenology of native catfishes in River Ganga and modeling species response to climatic variability for their conservation. - Int. J. Biometeorol. 63: 991–1004.

Sinclair, A. R. E. et al. 2000. What determines phenology and synchrony of ungulate breeding in Serengeti? -Ecology 81: 2100–2111.

Skinner, J. D. et al. 2002. Inherent seasonality in the breeding seasons of African mammals: evidence from captive breeding. - Trans. R. Soc. South Africa 57: 25–34.

Stearns, S. C. 1989. Trade-offs in life-history evolution. - Funct. Ecol. 3: 259–268.

van den Hoff, J. 2020. Environmental constraints on the breeding phenology of Giant Petrels Macronectes spp., with emphasis on Southern Giant Petrels M. giganteus. - Mar. Ornithol. 48: 33–40.

Visser, M. E. et al. 2010. Phenology, seasonal timing and circannual rhythms: towards a unified framework. - Philos. Trans. R. Soc. B Biol. Sci. 365: 3113–3127.

Zerbe, P. et al. 2012. Reproductive seasonality in captive wild ruminants: implications for biogeographical adaptation, photoperiodic control, and life history. - Biol. Rev. 87: 965–990.

