## Supporting information for "How to describe and measure phenology? An investigation on the diversity of metrics using phenology of births in large herbivores"

Supporting information 1: References of the articles from which the metrics describing phenology of births in large herbivores were extracted.


Aanes, R. and Andersen, R. 1996. The effects of sex, time of birth, and habitat on the vulnerability of roe deer fawns to red fox predation. - Can. J. Zool. 74: 1857–1865.

Adams, L. G. and Dale, B. W. 1998. Timing and synchrony of parturition in Alaskan caribou. - J. Mammal. 79: 287–294.

Bercovitch, F. B. and Berry, P. S. M. 2010. Reproductive life history of Thornicroft's giraffe in Zambia. - Afr. J. Ecol. 48: 535–538.

Berger, J. 1992. Facilitation of reproductive synchrony by gestation adjustment in gregarious mammals: a new hypothesis. - Ecology 73: 323–329.

Berger, J. and Cain, S. L. 1999. Reproductive synchrony in brucellosis-exposed bison in the southern Greater Yellowstone Ecosystem and in noninfected populations. - Conserv. Biol. 13: 357–366.

Bergerud, A. T. 1975. The reproductive season of Newfoundland caribou. - Can. J. Zool. 53: 1213–1221.

Bon, R. *et al.* 1993. Mating and lambing periods as related to age of female mouflon. - J. Mammal. 74: 752–757.

Bonnet, T. *et al.* 2019. The role of selection and evolution in changing parturition date in a red deer population. - PLoS Biol. 17: e3000493.

Bowyer, R. T. 1991. Timing of parturition and lactation in southern mule deer. - J. Mammal. 72: 138–145.

Bowyer, R. T. *et al.* 1998. Timing and synchrony of parturition in Alaskan moose: long-term versus proximal effects of climate. - J. Mammal. 79: 1332–1344.

Bunnell, F. L. 1980. Factors controlling lambing period of Dall's sheep. - Can. J. Zool. 58: 1027–1031.

Bunnell, F. L. 1982. The lambing period of mountain sheep: synthesis, hypotheses, and tests. - Can. J. Zool. 60: 1–14.

Calabrese, J. M. *et al.* 2018. Male rutting calls synchronize reproduction in Serengeti wildebeest. - Sci. Rep. 8: 10202.

Caughley, G. and Caughley, J. 1974. Estimating median date of birth. - J. Wildl. Manage.: 552–556.

English, A. K. *et al.* 2012. Reassessing the determinants of breeding synchrony in ungulates. - PLoS One 7: e41444.

Gaillard, J. M. *et al.* 1993. Timing and synchrony of births in roe deer. - J. Mammal. 74: 738–744.

Green, W. C. H. and Rothstein, A. 1993. Asynchronous parturition in bison: implications for the hider-follower dichotomy. - J. Mammal. 74: 920–925.

Guinness, F. E. *et al.* 1978. Calving times of red deer (Cervus elaphus) on Rhum. - J. Zool. 185: 105–114.

Hass, C. C. 1997. Seasonality of births in bighorn sheep. - J. Mammal. 78: 1251–1260.

Jarnemo, A. *et al.* 2004. Predation by red fox on European roe deer fawns in relation to age, sex, and birth date. - Can. J. Zool. 82: 416–422.

Johnson, D. S. *et al.* 2004. Estimating timing of life-history events with coarse data. - J. Mammal. 85: 932–939.

Lent, P. C. 1966. Calving and related social behavior in the barren-ground caribou. - Z. Tierpsychol. 23: 701–756.

Linnell, J. D. C. and Andersen, R. 1998. Timing and synchrony of birth in a hider species, the roe deer Capreolus capreolus. - J. Zool. 244: 497–504.

Loe, L. E. *et al.* 2005. Climate predictability and breeding phenology in red deer: timing and synchrony of rutting and calving in Norway and France. - J. Anim. Ecol. 74: 579–588.

McGinnes, B. S. and Downing, R. L. 1977. Factors affecting the peak of white-tailed deer fawning in Virginia. - J. Wildl. Manage.: 715–719.

Meng, X. *et al.* 2003. Timing and synchrony of parturition in Alpine musk deer (Moschus syfanicus). - FOLIA Zool. 52: 39–50.

Moe, S. R. *et al.* 2007. Trade-off between resource seasonality and predation risk explains reproductive chronology in impala. - J. Zool. 273: 237–243.

Nefdt, R. J. C. 1996. Reproductive seasonality in Kafue lechwe antelope. - J. Zool. 239: 155–166.

Ogutu, J. O. *et al.* 2010. Rainfall extremes explain interannual shifts in timing and synchrony of calving in topi and warthog. - Popul. Ecol. 52: 89–102.

Owen-Smith, N. and Ogutu, J. O. 2013. Controls over reproductive phenology among ungulates: allometry and tropical-temperate contrasts. - Ecography (Cop.). 36: 256–263.

Paoli, A. *et al.* 2018. Winter and spring climatic conditions influence timing and synchrony of calving in reindeer. - PLoS One 13: e0195603.

Paré, P. *et al.* 1996. Seasonal reproduction of captive Himalayan tahrs (Hemitragus jemlahicus) in relation to latitude. - J. Mammal. 77: 826–832.

Plard, F. *et al.* 2014. Mismatch between birth date and vegetation phenology slows the demography of roe deer. - PLoS Biol. 12: e1001828.

Post, E. 2003. Timing of reproduction in large mammals. - In: Phenology: an integrative environmental science. Springer, pp. 437–449.

Post, E. and Klein, D. R. 1999. Caribou calf production and seasonal range quality during a population decline. - J. Wildl. Manage.: 335–345.

Post, E. and Forchhammer, M. C. 2008. Climate change reduces reproductive success of an Arctic herbivore through trophic mismatch. - Philos. Trans. R. Soc. B Biol. Sci. 363: 2367–2373.

Post, E. *et al.* 2003. Synchrony between caribou calving and plant phenology in depredated and non-depredated populations. - Can. J. Zool. 81: 1709–1714.

Rachlow, J. L. and Bowyer, R. T. 1991. Interannual variation in timing and synchrony of parturition in Dall's sheep. - J. Mammal. 72: 487–492.

Renaud, L.-A. *et al.* 2019. Phenotypic plasticity in bighorn sheep reproductive phenology: from individual to population. - Behav. Ecol. Sociobiol. 73: 50.

Rutberg, A. T. 1984. Birth synchrony in American bison (Bison bison): response to predation or season? - J. Mammal. 65: 418–423.

Rutberg, A. T. 1987. Adaptive hypotheses of birth synchrony in ruminants: an interspecific test. - Am. Nat. 130: 692–710.

Ryan, S. J. *et al.* 2007. Ecological cues, gestation length, and birth timing in African buffalo (Syncerus caffer). - Behav. Ecol. 18: 635–644.

Sigouin, D. *et al.* 1997. PERIODS OF MOOSE. - Alces 33: 85–95.

Sinclair, A. R. E. *et al.* 2000. What determines phenology and synchrony of ungulate breeding in Serengeti? - Ecology 81: 2100–2111.

Skinner, J. D. *et al.* 2002. Inherent seasonality in the breeding seasons of African mammals: evidence from captive breeding. - Trans. R. Soc. South Africa 57: 25–34.

Whiting, J. C. *et al.* 2012. Timing and synchrony of births in bighorn sheep: implications for reintroduction and conservation. - Wildl. Res. 39: 565–572.

Zerbe, P. *et al.* 2012. Reproductive seasonality in captive wild ruminants: implications for biogeographical adaptation, photoperiodic control, and life history. - Biol. Rev. 87: 965–990.

Supporting information 2: Parameters used in the simulations of phenology of births and default parameters used to implement the phenology metrics.

Table S2-1: Ranges of the four parameters varying in the simulated phenology of births: mean birth date for a given year, standard deviation of the birth distribution for a given year, range over which the mean birth date can vary across years, range over which the standard deviation can vary across years.

| **Characteristic of phenology** | **Observed parameter** | **Mini-mum value** | **Maxi-mum value** | **Incre-ment** | **Number of values** | **Number of repetitions** | **Total number of simulations** |
|---|---|---|---|---|---|---|---|
| Timing | mean | 85 | 283 | 4 | 50 | 50 | 2500 |
| Synchrony | standard deviation (sd) | 1 | 75 | 1.5 | 50 | 50 | 2500 |
| Rhythmicity | Δmean | 10 | 100 | 10 | 10 | 250 | 2500 |
| Regularity | Δsd | 3 | 40 | 4 | 10 | 250 | 2500 |

Table S2-2: Default parameters for the metrics, selected according to the most common practice in the literature of phenology of births in large herbivores.

| **Parameter** | **Value selected** | **Reference** |
|---|---|---|
| Minimum percentage of births to consider that births are synchronous | 80 % | Rutberg 1987 |
| First quantile of the distribution of births (to evaluate births synchrony) | 25 % | Gaillard *et al.* 1993 |
| Last quantile of the distribution of births (to evaluate births synchrony) | 75 % | Gaillard *et al.* 1993 |
| Minimum percentage of the total number of births occurring within a year that should happen in a given time unit (*e.g.* a month) to count this time unit as a time unit during which births occur ("*nbtu*" metric) | 1 % | Moe *et al.* 2007 |
| Birth should be accounted for only if they are distributed in consecutive time units (*e.g.* months) | FALSE | / |
| Number of repetitions for the simulation procedures | 1000 | / |
| Transformation of the data | NONE | / |
| Confidence intervals | 95 % | / |
| Period to consider that births are synchronous | Mean theoretical standard deviation of all the 2500 patterns simulated for which standard deviation varies | / |

Because the implementation of a few metrics relied on the validation or invalidation of *a priori* condition (*e.g.* "at least a certain percentage of births occurs during a given period" to determine whether or not births are synchronous), we selected the most commonly used setting for that particular metric according to what we encountered in the literature or settings as general as possible to suit a large range of phenology of births.

When more than one value was possibly returned by the function coding for a metric, we retained only one value according to a predefined rule. For metrics returning the dates of the peaks in the distribution of births (*n* = 2 metrics), we only kept the date of the first peak if more than one peak was detected. When the metric was a boolean (true/false) variable based on the significance of a statistical test (*n* = 9 metrics), we used the value of the test statistics as output metric, thereby allowing us to investigate how the statistics was influenced by the value of phenology parameters. When the metric was boolean but not based on a statistical test (*n* = 4 metrics), we coded its value as binary variable taking a "1" when the test was significant or "0" when not significant at the *alpha* = 0.05 level. When the metric could take both positive and negative values representing the direction of a deviation such as the skewness (*n* = 2 metrics), we used the absolute value as we were interested in the amplitude and not in the direction of the deviation. One metric ("*khi2*" metric) could not be used for some simulations (1.5 % of the simulations) because the range of birth dates was sometimes too small to run the function associated with this metric.

Supporting information 3: Detailed description of the metrics extracted from phenology of births in large herbivore literature.

The description of each metric (*e.g.* characteristic of phenology associated, number of years it requires to be implemented, null hypothesis tested if this is the case) are provided. The details of the score obtained by each metric for each criterion listed in Table 1 are provided too.

Table S3-1: General characteristics of the metrics.

| **Metric**[1] | **Complete name** | **Theoretical phenology characteristic associated**[2] | **Observed phenology characteristic associated**[3] | **Appli-cation period**[4] | **Unit**[5] | **Reference** |
|---|---|---|---|---|---|---|
| bart | bartlett | regularity | regularity | several cycles | boolean | Hass 1997 |
| bgper | beginning period | timing | timing+ | one cycle | date | Lent 1966 |
| bgthper | beginning period threshold | timing | timing+ | one cycle | date | Post and Forchhammer 2008 |
| centre | centre | timing | timing | one cycle | date | Sigouin *et al.* 1997 |
| cmano | comparison mean anova | rhythmicity | rhythmicity | several cycles | boolean | Gaillard *et al.* 1993 |
| compmean | comparison mean ci | rhythmicity | rhythmicity | two cycles | boolean | Whiting *et al.* 2012 |
| comppeaksig | comparison peak sigmoid ci | rhythmicity | none | two cycles | boolean | Moe *et al.* 2007 |
| diffbgper | difference beginning period | rhythmicity | rhythmicity+ | two cycles | duration | Guinness *et al.* 1978 |
| diffmean | difference mean linear | rhythmicity | rhythmicity | several cycles | duration | Paoli *et al.* 2018 |
| diffmed | difference median | rhythmicity | rhythmicity+ | two cycles | duration | Berger 1992 |
| diffmima | difference min max proportion births | synchrony | synchrony | one cycle | count | Owen-Smith and Ogutu 2013 |
| diffpeak | difference peak | rhythmicity | rhythmicity+ | two cycles | duration | Guinness *et al.* 1978 |
| diffperiod | difference period | regularity | regularity | two cycles | duration | Berger and Cain 1999 |
| diffslin | difference synchrony linear | regularity | regularity+ | several cycles | duration | Paoli *et al.* 2018 |
| interq | interquantiles period quantiles | synchrony | synchrony | one cycle | duration | Gaillard *et al.* 1993 |
| khi2 | khi2 proportion births median | regularity | regularity | several cycles | boolean | Adams and Dale 1998 |
| kolmogau | kolmogorov smirnov gaussian | synchrony | synchrony | one cycle | boolean | Linnell and Andersen 1998 |

| | | | | | | |
|---|---|---|---|---|---|---|
| kolmomult | kolmogorov smirnov multi-year | rhythmicity - regularity | regularity+ | two cycles | boolean | Green and Rothstein 1993 |
| kolmouni | kolmogorov smirnov uniform | synchrony | synchrony | one cycle | boolean | Sinclair *et al.* 2000 |
| maxprop | max proportion births period given | synchrony | synchrony | one cycle | count | Owen-Smith and Ogutu 2013 |
| mean | mean | timing | timing | one cycle | date | McGinnes and Downing 1977 |
| meanlin | mean linear random | timing | timing | several cycles | date | Loe *et al.* 2005 |
| meanmult | mean mean multi-year | timing | timing | several cycles | date | Jarnemo *et al.* 2004 |
| meanvl | mean vector length | synchrony | synchrony | one cycle | unitless value | Paré *et al.* 1996 |
| meanvo | mean vector orientation | timing | timing | one cycle | date (radian) | Paré *et al.* 1996 |
| med | median | timing | timing | one cycle | date | Bergerud 1975 |
| medprob | median probit | timing | timing | one cycle | date | Caughley and Caughley 1974 |
| minper | min period proportion births given | synchrony | synchrony | one cycle | duration | Skinner *et al.* 2002 |
| minprop | min proportion births period given | synchrony | synchrony | one cycle | count | Owen-Smith and Ogutu 2013 |
| mode | mode | timing | timing | one cycle | date | Bergerud 1975 |
| mood | mood | rhythmicity | rhythmicity | two cycles | boolean | Berger and Cain 1999 |
| nbtu | number time unit minimal births | synchrony | synchrony | one cycle | duration | Moe *et al.* 2007 |
| peaksig | peak sigmoid | timing | timing | one cycle | date | Moe *et al.* 2007 |
| per | period | synchrony | synchrony | one cycle | duration | Lent 1966 |
| pergau | period gaussian | synchrony | synchrony | one cycle | duration | Paoli *et al.* 2018 |
| perhdr | period high density region | synchrony | synchrony | one cycle | duration | Calabrese *et al.* 2018 |
| permean | period mean multi-year | synchrony | synchrony | several cycles | duration | Green and Rothstein 1993 |
| pielou | pielou | synchrony | synchrony | one cycle | unitless value | Sinclair *et al.* 2000 |
| propmed | proportion births around median | synchrony | synchrony | one cycle | count | Adams and Dale 1998 |
| propmode | proportion births around mode | synchrony | synchrony | one cycle | count | Green and Rothstein 1993 |
| rayleigh | rayleigh | synchrony | none | one cycle | boolean | Paré *et al.* 1996 |
| rutberg | rutberg | synchrony | synchrony | one cycle | duration | Rutberg 1984 |
| sd | standard deviation | synchrony | synchrony | one cycle | duration | Bowyer 1991 |
| sdprob | standard deviation probit | synchrony | synchrony | one cycle | duration | Caughley and Caughley 1974 |

| | | | | | | |
|---|---|---|---|---|---|---|
| skew | skewness variance | synchrony | none | one cycle | unitless | Bunnell 1980 |
| skinner | skinner | synchrony | synchrony | one cycle | boolean | Skinner *et al.* 2002 |
| slpcomp | slope comparison | regularity | synchrony | several cycles | boolean | Bowyer *et al.* 1998 |
| var | variance | synchrony | synchrony | one cycle | duration | Hass 1997 |
| varcor | variance corrected | synchrony | synchrony | one cycle | duration | Johnson *et al.* 2004 |
| varlin | variance mutli-year | rhythmicity | rhythmicity+ | several cycles | duration | Loe *et al.* 2005 |
| watson | watson williams | rhythmicity | rhythmicity | two cycles | boolean | Paré *et al.* 1996 |
| zerbe | zerbe | synchrony | synchrony | one cycle | duration | Zerbe *et al.* 2012 |

[1] short name of the metric as used in the article; [2] characteristic of phenology (timing, synchrony, rhythmicity or regularity) the metric is theoretically supposed to measure; [3] characteristic of phenology (timing, synchrony, rhythmicity or regularity) the metric effectively measures according to our analyses (+: when the metric also vary according to other characteristics of phenology than the one expected); [4] is the metric applicable to one year, two years or several years; [5] in which unit is the metric (days, number of births, a boolean as true or false, *etc.*).

Table S3-2: Detailed description of the metrics.

| Metric | Description[1] | Tested hypothesis[2] |
|---|---|---|
| bart | compare variances of birth distributions | Variances of the birth distributions are similar |
| bgper | find first birth date | |
| bgthper | find first birth date when at least x percent of births have occurred | |
| centre | find central date between first and last birth dates | |
| cmano | compare mean birth dates between several reproductive cycles thanks to one way anova | Median birth dates are similar |
| compmean | compare mean birth dates between two reproductive cycles | Confidence intervals of mean birth dates overlap |
| comppeaksig | compare date of inflection point of birth distribution between two reproductive cycles | Confidence intervals of inflection points overlap |
| diffbgper | evaluate duration between first birth dates of two reproductive cycles | |
| diffmean | evaluate slope coefficient of linear model describing distribution of mean birth dates of several reproductive cycles | |
| diffmed | evaluate duration between median birth dates of two reproductive cycles | |
| diffmima | evaluate difference between "maxprop" and "minprop" metrics | |
| diffpeak | evaluate duration between mode birth dates of two reproductive cycles | |
| diffperiod | evaluate difference of duration between birth period duration (period between first and last birth dates) of two reproductive cycles | |
| diffslin | evaluate slope coefficient of linear model describing distribution of "pergau" metric of several reproductive cycles | |
| interq | find period gathering x percent of births based on quantiles | |
| khi2 | compare distribution of proportion of births around median birth date of several reproductive cycles to a uniform distribution | Distribution of birth proportions around median birth date follows a uniform distribution |
| kolmogau | compare birth distribution to a gaussian distribution | Birth distribution follows a normal distribution |
| kolmomult | compare birth distribution between two reproductive cycles | Birth distribution similar for both reproduction cycles |
| kolmouni | compare birth distribution to a uniform distribution | Birth distribution follows a uniform distribution |
| maxprop | evaluate maximum proportion of births for a given duration | |
| mean | find mean birth date | |
| meanlin | find mean birth date thanks to linear model with random effects | |
| meanmult | calculate mean of mean birth dates | |
| meanvl | evaluate mean vector length (circular statistics) | |
| meanvo | evaluate mean vector orientation (circular statistics) | |
| med | find median birth date | |
| medprob | evaluate median birth date thanks to probit analysis | |

| | | |
|---|---|---|
| minper | evaluate shortest period gathering x percent of births | |
| minprop | evaluate proportion of births occurring during the consecutive period gathering the less births | |
| mode | find mode birth date | |
| mood | compare median birth dates between two reproductive cycles | Median birth dates are similar |
| nbtu | find duration gathering at least x percent of the births | |
| peaksig | find inflection point based on logistic regression describing cumulative births | |
| per | calculate duration between first and last birth | |
| pergau | calculate 2*2*standard deviation of birth distribution | |
| perhdr | evaluate duration gathering x percent of births thanks to high density regions | |
| permean | evaluate mean duration between first and last births | |
| pielou | evaluate evenness index | |
| propmed | evaluate proportion of births occurring around median birth date | |
| propmode | evaluate proportion of births occurring around mode birth date | |
| rayleigh | compare birth distribution to a uniform distribution (alternative hypothesis: unimodal distribution) | Birth distribution follows a random distribution |
| rutberg | evaluate shortest period gathering at least x percent of births since first birth | |
| sd | calculate standard deviation of birth distribution | |
| sdprob | calculate standard deviation of birth distribution based on probit analysis | |
| skew | evaluate skewness of birth distribution | |
| skinner | evaluate presence or absence of a period of a given duration gathering x percent of births | At least one period of x days gathering y percents of births |
| slpcomp | compare slope coefficients of linear models describing birth distributions after transformation | Slopes of the linear regressions of the number of births according to time are similar |
| var | calculate variance of birth distribution | |
| varcor | calculate variance of birth distribution corrected by the Sheppard method | |
| varlin | calculate inter-reproductive cycles variance thanks to linear model with random effects | |
| watson | compare mean birth dates between two reproductive cycles | Mean birth dates are similar |
| zerbe | find shortest period gathering x percent of births around mode birth date | |

[1] brief description of the metric and what it evaluates; [2] null hypotheses tested (when possible).

Table S3-3: Scores of the metrics.

| Metric | Theoretical phenology characteristic associated | Goodness[1] | Monotony[1] | Saturation[1] | Strength[1] | Normality (assumption) [1] | Normality (robustness) [1] | Origin[1] | Linearity[1] | Unicity[1] | Score[2] |
|---|---|---|---|---|---|---|---|---|---|---|---|
| bart | regularity | T | T | T | medium | F | T | F | 2 | T | 6 |
| bgper | timing | T | T | T | high | T | | F | 1 | T | 7 |
| bgthper | timing | T | T | T | high | T | | F | 1 | T | 7 |
| centre | timing | T | T | T | high | T | | F | 1 | T | 7 |
| cmano | rhythmicity | T | T | F | | | | | | | 2 |
| compmean | rhythmicity | T | T | F | | | | | | | 2 |
| comppeaksig | rhythmicity | F | | | | | | | | | 0 |
| diffbgper | rhythmicity | T | T | T | medium | T | | F | 1 | T | 6.5 |
| diffmean | rhythmicity | T | T | F | | | | | | | 2 |
| diffmed | rhythmicity | T | T | T | high | T | | F | 1 | T | 7 |
| diffmima | synchrony | T | T | F | | | | | | | 2 |
| diffpeak | rhythmicity | T | T | T | medium | T | | F | 1 | F | 5.5 |
| diffperiod | regularity | T | T | T | medium | T | | F | 1 | T | 6.5 |
| diffslin | regularity | T | T | F | | | | | | | 2 |
| interq | synchrony | T | T | T | high | T | | F | 1 | T | 7 |
| khi2 | regularity | T | T | T | medium | T | | F | 3 | T | 6 |
| kolmogau | synchrony | T | T | F | | | | | | | 2 |
| kolmomult | rhythmicity - **regularity** | T | T | T | medium | T | | T | 1 | T | 7.5 |
| kolmouni | synchrony | T | T | T | high | T | | T | 3 | T | 7.5 |
| maxprop | synchrony | T | T | F | | | | | | | 2 |
| mean | timing | T | T | T | high | T | | F | 1 | T | 7 |
| meanlin | timing | T | T | T | high | T | | F | 1 | T | 7 |
| meanmult | timing | T | T | T | high | T | | F | 1 | T | 7 |
| meanvl | synchrony | T | T | T | high | T | | T | 3 | T | 7.5 |
| meanvo | timing | T | T | T | high | T | | T | 1 | T | 8 |
| med | timing | T | T | T | high | T | | F | 1 | T | 7 |
| medprob | timing | T | T | T | high | F | T | F | 1 | T | 7 |
| minper | synchrony | T | T | T | high | T | | F | 1 | F | 6 |
| minprop | synchrony | T | T | F | | | | | | | 2 |
| mode | timing | T | T | T | medium | T | | T | 1 | F | 6.5 |
| mood | rhythmicity | T | T | T | high | T | | F | 3 | T | 6.5 |
| nbtu | synchrony | T | F | | | | | | | | 1 |
| peaksig | timing | T | T | T | high | F | T | F | 1 | T | 7 |
| per | synchrony | T | T | F | | | | | | | 2 |
| pergau | synchrony | T | T | T | high | F | T | F | 1 | T | 7 |
| perhdr | synchrony | T | T | T | high | T | | F | 1 | F | 6 |
| permean | synchrony | T | T | F | | | | | | | 2 |

| | | | | | | | | | | | |
|---|---|---|---|---|---|---|---|---|---|---|---|
| pielou | synchrony | T | T | T | high | T | | T | 3 | T | 7.5 |
| propmed | synchrony | T | T | F | | | | | | | 2 |
| propmode | synchrony | T | T | F | | | | | | | 2 |
| rayleigh | synchrony | F | | | | | | | | | 0 |
| rutberg | synchrony | T | T | T | medium | T | | F | 1 | T | 6.5 |
| sd | synchrony | T | T | T | high | T | | F | 1 | T | 7 |
| sdprob | synchrony | T | T | T | high | F | T | F | 1 | T | 7 |
| skew | synchrony | F | | | | | | | | | 0 |
| skinner | synchrony | T | T | F | | | | | | | 2 |
| slpcomp | regularity | F | | | | | | | | | 0 |
| var | synchrony | T | T | T | high | T | | F | 3 | T | 6.5 |
| varcor | synchrony | T | T | T | high | T | | F | 3 | T | 6.5 |
| varlin | rhythmicity | T | T | T | high | T | | F | 3 | T | 6.5 |
| watson | rhythmicity | T | T | T | high | F | T | F | 3 | T | 6.5 |
| zerbe | synchrony | T | T | T | high | T | | F | 1 | F | 6 |

[1] *cf.* Table 1 in the main text; [2] total score obtained by the metric according to the eight criteria.

Supporting information 4: Evaluation of phenology metrics in scenarios of non-normal distributions of births.

**Introduction**

In the main text, we chose to base our simulations on a normal distribution of births for biological and methodological considerations (see Materials and Methods section). Normally distributed dates of birth were reported for roe deer *Capreolus capreolus* (Gaillard *et al.* 1993) and wildebeest *Connochaetes taurinus* for instance (Sinclair *et al.* 2000). However, the distribution of births of some species is not necessarily normal. Here, we replicated our analyses on the metrics with additional distributions of births previously reported in the literature. We identified four scenarios (Fig. S4-1):

1) a **skewed normal distribution**, which would better approximate distributions of births characterized by numerous early births and fewer late births. Such asymmetric distributions of dates of birth were documented for warthog *Phacochoerus aethiopicus* (Sinclair *et al.* 2000), or bighorn sheep *Ovis canadensis* (Festa-Bianchet 1988) for instance;

2) a **bimodal distribution with two close peaks**, which would better approximate distributions of births characterized by a main peak closely followed by a second one, often smaller. Such distributions could arise from second attempts to breed for females whose first attempt failed (*e.g.* because fertilization failed at the first oestrus). Examples of species displaying such a bimodal distribution of dates of birth are the impala *Aepyceros melampus* (Anderson 1975) or the red deer *Cervus elaphus* (Guinness *et al.* 1978);

3) a **Cauchy distribution**, which represents a bell-shape distribution of births with fat tails. Even if such distribution looks like a normal distribution at first glance, it is however characterized by the existence of few births occurring all year long in addition to the main delimited peak. The distribution of births of zebra *Equus burchelli böhmi* (Leuthold and

Leuthold 1975) or Grant's gazelle *Gazella granti* (Sinclair *et al.* 2000) match with this theoretical distribution;

4) a **random distribution** whereby births can occur anytime in the year because conditions are always favorable, for instance. Giraffe *Giraffa camelopardalis* (Sinclair *et al.* 2000) and waterbuck *Kobus ellipsiprymnus* (Leuthold and Leuthold 1975) can give birth all year round.

### Materials and Methods

Here, we evaluated the behaviour of each metric presented in the main text and in Supporting information 3, according to the four characteristics of phenology (timing, synchrony, rhythmicity and regularity). We replicated the methodology described in "Materials and Methods" section of the main text, Steps 2 to 4. Each simulated phenology of births was generated following one of the four distributions presented above by distributing approx. 1000 births within a year of 365 days, replicated over 10 years (Step 2). According to the distribution, we fixed specific parameters and we changed our four parameters of interest independently (Table S4-1) to illustrate variations of: *i*) the timing (*i.e.* mean day of birth for a given year), *ii*) the synchrony (*i.e.* standard deviation of the distribution of births for a given year), *iii*) the rhythmicity (*i.e.* size of the range over which the mean birth date can vary across years), and *iv*) the regularity (*i.e.* size of the range over which the standard deviation can vary across years). Each of those parameters varied in a range from a minimum to a maximum value and was incremented with a constant step. We then computed the metrics from each simulated phenology following Step 3. We finally produced the global correlation matrix between all pairs of metrics, using Pearson correlations, for each distribution (Step 4).

We compared the results obtained using the above-described distributions with those obtained when using a normal distribution. To do so, for each metric, we (1) extracted the correlation coefficients between this metric and the others while using a normal distribution, (2) did the same with correlation values obtained when using other distributions, and (3) fitted a linear model between values obtained in (2) and those obtained in (1), thereby testing to what extent values obtained using non-normal distributions could be predicted from those obtained using normal distributions. We used the coefficient of determination $R^2$ as a measure of the fit. A high $R^2$ indicated that the correlation between metrics was not greatly affected by the choice of the distribution of births.

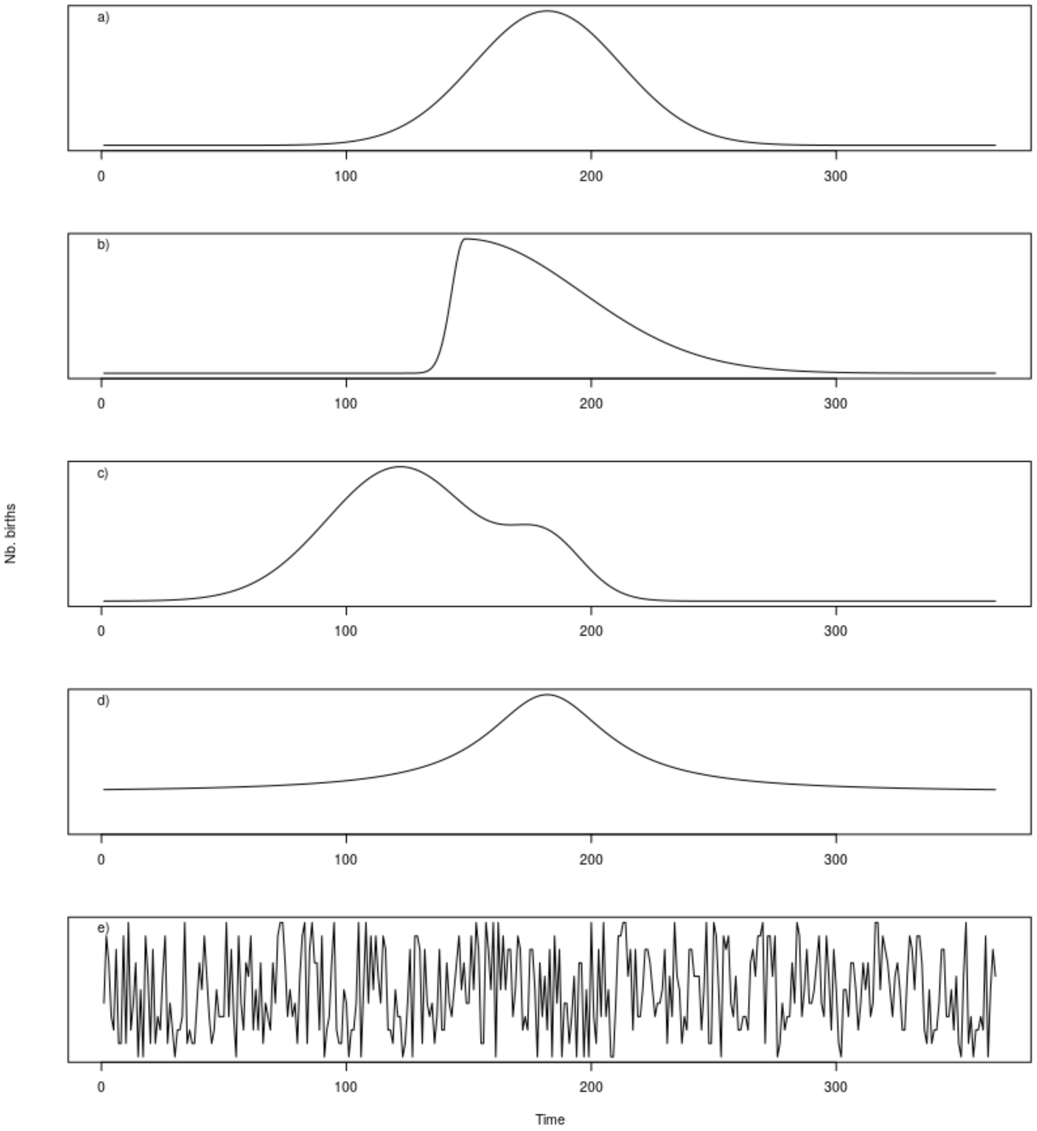


Figure S4-1: theoretical distributions that could illustrate phenology of births of large herbivore species in *natura*: a) normal distribution, b) skewed normal distribution, c) bimodal distribution with two close peaks, d) Cauchy distribution, e) random distribution.

Table S4-1: Values of the parameters used in the simulated phenology of births following the four scenarios (skew normal distribution, bimodal distribution with two close peaks, Cauchy distribution, random distribution). For the variable parameters (mean, standard deviation, range of variation of both parameters), 10 different values were used, with 10 repetitions each time, leading to 100 simulations per unique combination of parameters.

| Distribution | Characteristic of phenology | Observed parameter | Default value | Minimum value | Maximum value | Incre-ment |
|---|---|---|---|---|---|---|
| Skewed normal | Timing | mean | 182 | 90 | 270 | 20 |
| | Synchrony | standard deviation (sd) | 30 | 2 | 74 | 8 |
| | Rhythmicity | Δmean | 0 | 10 | 100 | 10 |
| | Regularity | Δsd | 0 | 5 | 41 | 4 |
| | Skewness | skew | 3.5 | | | |
| Bimodal | Timing | mean | 122 | 90 | 225 | 15 |
| | Synchrony | sd | 30 | 10 | 37 | 3 |
| | Rhythmicity | Δmean | 0 | 5 | 50 | 5 |
| | Regularity | Δsd | 0 | 3 | 30 | 3 |
| | Distance between main and second peak | Δmean/mean2 | 60 | | | |
| | Synchrony second peak | sd2 | 15 | | | |
| | Proportion of births in main peak | pb1 | 0.75 | | | |
| Cauchy | Timing | mean | 182 | 90 | 270 | 20 |
| | Synchrony | sd | 30 | 2 | 74 | 8 |
| | Rhythmicity | Δmean | 0 | 10 | 100 | 10 |
| | Regularity | Δsd | 0 | 5 | 41 | 4 |
| Random | Maximum number of births per day | $max_{nb}$ | 5 | | | |

**Results**

Most of the relationships between the correlation coefficients based on the skewed normal, bimodal or Cauchy distributions and the correlation coefficients based on the normal distribution showed a $R^2 > 0.75$ (96 %, 71 % and 80 % of the metrics for the skewed normal, bimodal and Cauchy distributions respectively, Table S4-2). Similarly, less than 6 % of the relationships showed a $R^2 < 0.25$ for those three distributions. Although the results were generally congruent with those we reported when using the normal distribution as the baseline distribution of births (compare Fig. 2 in the main text and Fig. S4-2, a), b) and c)), some metrics measuring the same characteristic of phenology in the context of a normal distribution were less correlated when applied to non-normal distribution of dates of birth. The metric "*diffpeak*" evaluates the duration between the mode birth dates of two years. Depending on whether the mode is located in the first or the second peak in each year, the difference returned by "*diffpeak*" can vary in the bimodal distribution. Also, "*rutberg*", "*diffperiod*", "*bgper*" and "*bgthper*" suffered from the absence of real break in the Cauchy distribution. All these metrics rely on the detection of the beginning of a definite period of births so the absence of any period with no birth limits their relevance. To the contrary, some metrics did better when applied to the three asymmetrical patterns of births. This was the case for "*skew*", which measures the skewness of the distribution, and "*kolmogau*", which compares the distribution of births to a normal distribution. The two metrics correlated with the other synchrony metrics for those three distributions better than for the normal distribution. Indeed, "*skew*" detects the skewness of a distribution, a feature existing in the three distributions but not in the normal one. "*kolmogau*" detects patterns departing from the normal distribution in the three other scenarios.

To the contrary, the random distribution of births produced very different results from all the other distributions (Fig. S4-2, d)). None of the relationships between the correlation coefficients for the random distribution and the correlation coefficients based on the normal distribution had a $R^2 > 0.75$, and 67 % had a $R^2 < 0.50$ (Table S4-2). Some synchrony and timing metrics remained highly correlated, mainly because births were quite consistent through the year, such as the mean date of births ("*mean*") or the variance of the distribution of births ("*var*"), but it is statistically and biologically meaningless to characterize such distribution by its mean or variance though.

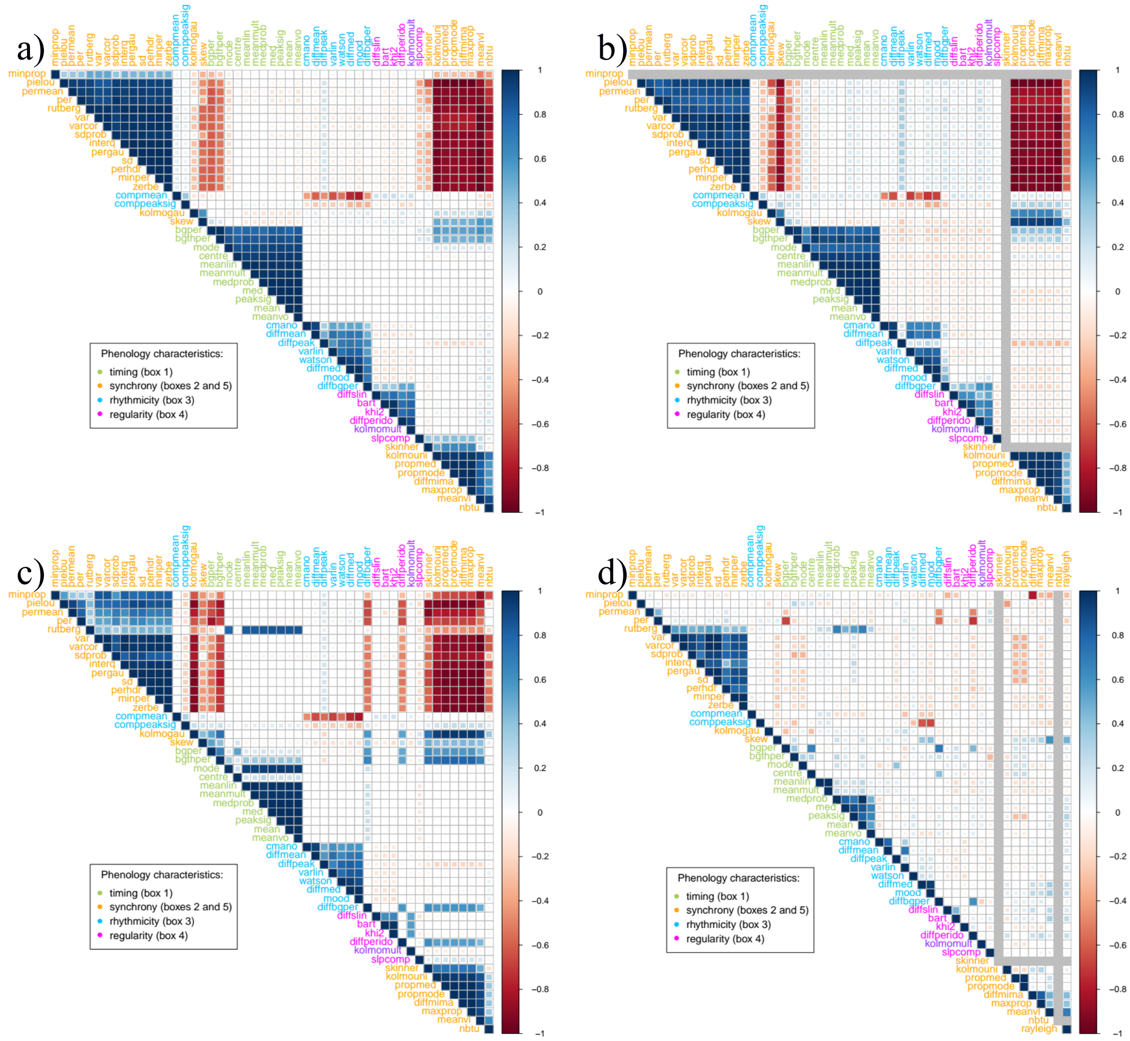


Figure S4-2: Correlation matrices between all pairs of metrics using Pearson correlations based on four scenarios: a) skewed normal distribution, b) bimodal distribution with two close peaks, c) Cauchy distribution, d) random distribution. It was not possible to classify "*kolmomult*" *a priori* in rhythmicity or regularity metrics, as it compares the complete distribution of births between two years. *Green* = timing metrics, *orange* = synchrony metrics, *blue* = rhythmicity metrics, *pink* = regularity metrics. When the value of a given metric was constant for a given distribution, it was not possible to assess the coefficients of correlation (reported in grey in the matrices).

Table S4-2: Comparison of the coefficients of correlation between all pairs of metrics based on the normal distribution and those based on the other distributions (skew normal distribution, distribution with two close peaks, Cauchy distribution and random distribution), using the coefficient of determination of the linear relationship. When the value of a given metric was constant either for the normal or the non-normal distributions, it was not possible to assess the coefficients of correlation, so the linear relationship was not explored (reported as “not applicable” in the table).

| **Metric** | **Skewed normal** | **Bimodal** | **Cauchy** | **Random** |
|---|---|---|---|---|
| bart | 0.98 | 0.7 | 0.82 | 0.42 |
| bgper | 0.98 | 0.85 | 0.7 | 0.24 |
| bgthper | 0.98 | 0.8 | 0.74 | 0.37 |
| centre | 0.99 | 0.9 | 0.56 | 0.09 |
| cmano | 0.99 | 0.85 | 0.94 | 0.56 |
| compmean | 0.98 | 0.67 | 0.9 | 0.45 |
| comppeaksig | 0.79 | 0.26 | 0.36 | 0.68 |
| diffbgper | 0.89 | 0.46 | 0.11 | 0.16 |
| diffmean | 0.98 | 0.8 | 0.93 | 0.56 |
| diffmed | 0.98 | 0.72 | 0.9 | 0.48 |
| diffmima | 0.98 | 0.89 | 0.89 | 0.34 |
| diffpeak | 0.97 | 0.59 | 0.92 | 0.19 |
| diffperido | 0.96 | 0.57 | 0.12 | 0.29 |
| diffslin | 0.98 | 0.71 | 0.78 | 0.37 |
| interq | 0.98 | 0.88 | 0.92 | 0.54 |
| khi2 | 0.97 | 0.72 | 0.76 | 0.26 |
| kolmogau | 0.46 | 0.2 | 0.1 | 0.64 |
| kolmomult | 0.97 | 0.73 | 0.59 | 0.16 |
| kolmouni | 0.98 | 0.89 | 0.9 | 0.11 |
| maxprop | 0.98 | 0.89 | 0.89 | 0.48 |
| mean | 0.99 | 0.95 | 0.78 | 0.51 |
| meanlin | 0.99 | 0.95 | 0.78 | 0.28 |
| meanmult | 0.99 | 0.95 | 0.78 | 0.28 |
| meanvl | 0.97 | 0.89 | 0.91 | 0.38 |
| meanvo | 0.97 | 0.94 | 0.78 | 0.47 |
| med | 0.98 | 0.91 | 0.78 | 0.54 |
| medprob | 0.99 | 0.97 | 0.79 | 0.49 |
| minper | 0.98 | 0.89 | 0.91 | 0.59 |
| minprop | 0.93 | Not applicable | 0.92 | 0.15 |
| mode | 0.94 | 0.87 | 0.78 | 0.2 |
| mood | 0.98 | 0.68 | 0.9 | 0.46 |
| nbtu | 0.97 | 0.86 | 0.79 | Not applicable |
| peaksig | 0.98 | 0.93 | 0.78 | 0.39 |
| per | 0.98 | 0.9 | 0.78 | 0.26 |
| pergau | 0.98 | 0.9 | 0.91 | 0.59 |
| perhdr | 0.98 | 0.88 | 0.91 | 0.57 |

| | | | | |
|---|---|---|---|---|
| permean | 0.98 | 0.89 | 0.81 | 0.19 |
| pielou | 0.98 | 0.89 | 0.88 | 0.09 |
| propmed | 0.98 | 0.89 | 0.89 | 0.37 |
| propmode | 0.98 | 0.89 | 0.89 | 0.37 |
| rayleigh | Not applicable | Not applicable | Not applicable | Not applicable |
| rutberg | 0.97 | 0.89 | 0.41 | 0.29 |
| varlin | 0.99 | 0.77 | 0.92 | 0.54 |
| skew | 0.34 | 0.23 | 0.35 | 0.48 |
| skinner | 0.98 | Not applicable | 0.76 | Not applicable |
| slpcomp | 0.95 | 0.22 | 0.83 | 0.22 |
| sd | 0.98 | 0.9 | 0.91 | 0.59 |
| sdprob | 0.98 | 0.9 | 0.9 | 0.56 |
| var | 0.97 | 0.89 | 0.92 | 0.61 |
| varcor | 0.97 | 0.89 | 0.92 | 0.61 |
| watson | 0.98 | 0.82 | 0.91 | 0.29 |
| zerbe | 0.98 | 0.89 | 0.91 | 0.59 |

Supporting information 5: Variation of each phenological metric according to the variation of the four parameters of phenology, within and between years, in the simulations of phenology of births. *mean*: variation of each metric according to the variation of the mean birth date in a give year; *sd*: variation of each metric according to the variation of the standard deviation of the distribution of births for a given year; *Δmean*: variation of each metric according to the variation of the size of the range over which the mean birth date can vary across years; *Δsd*: variation of each metric according to the variation of the size of the range over which the standard deviation of the distribution of births can vary across years.

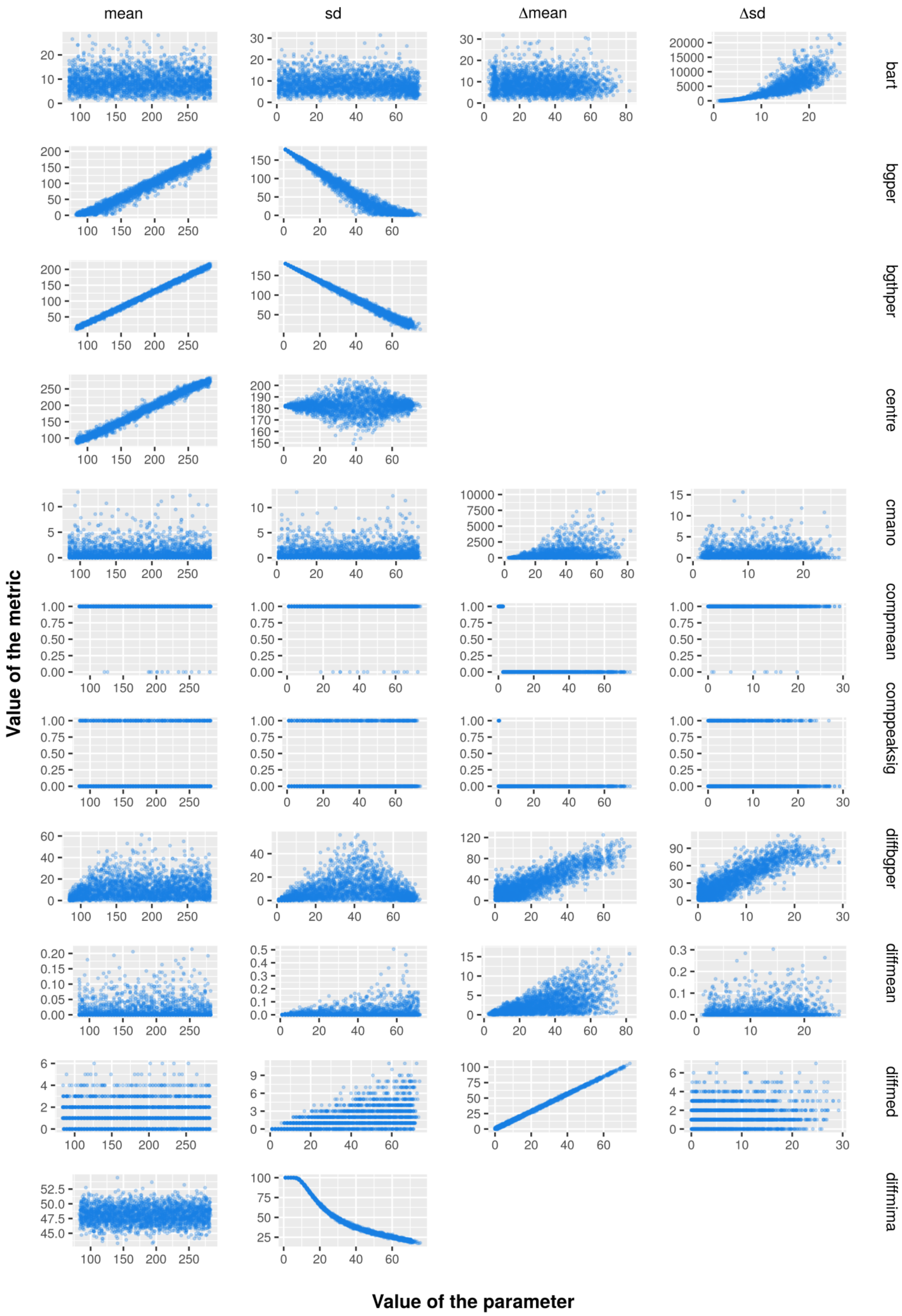
mean
sd
Δmean
Δsd
bart
bgper
bgthper
centre
cmano
compmean
comppeaksig
diffbgper
diffmean
diffmed
diffmima
Value of the metric
Value of the parameter

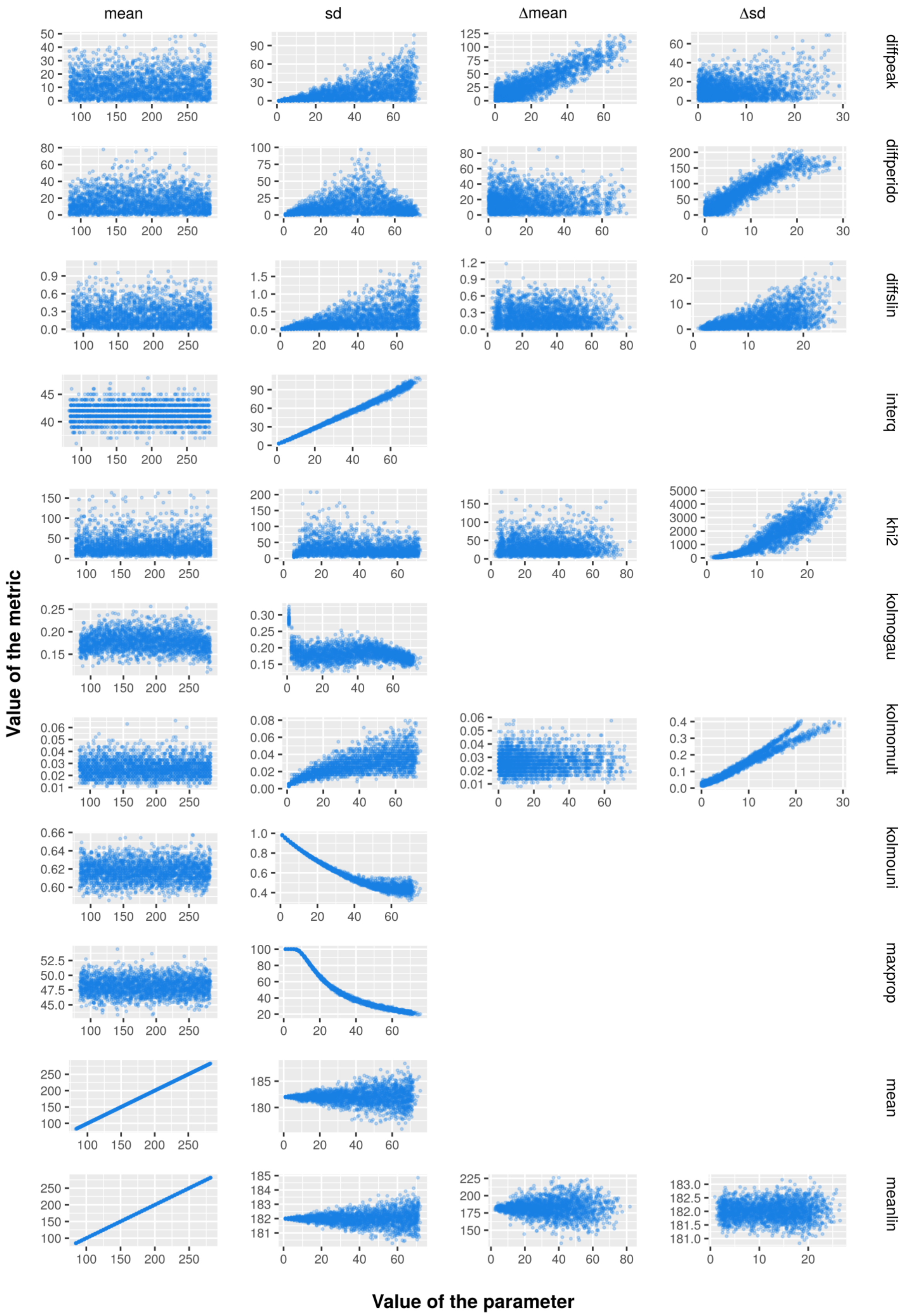
mean
sd
Δmean
Δsd
diffpeak
diffperido
diffslin
interq
khi2
kolmogau
kolmomult
kolmouni
maxprop
mean
meanlin
Value of the metric
Value of the parameter

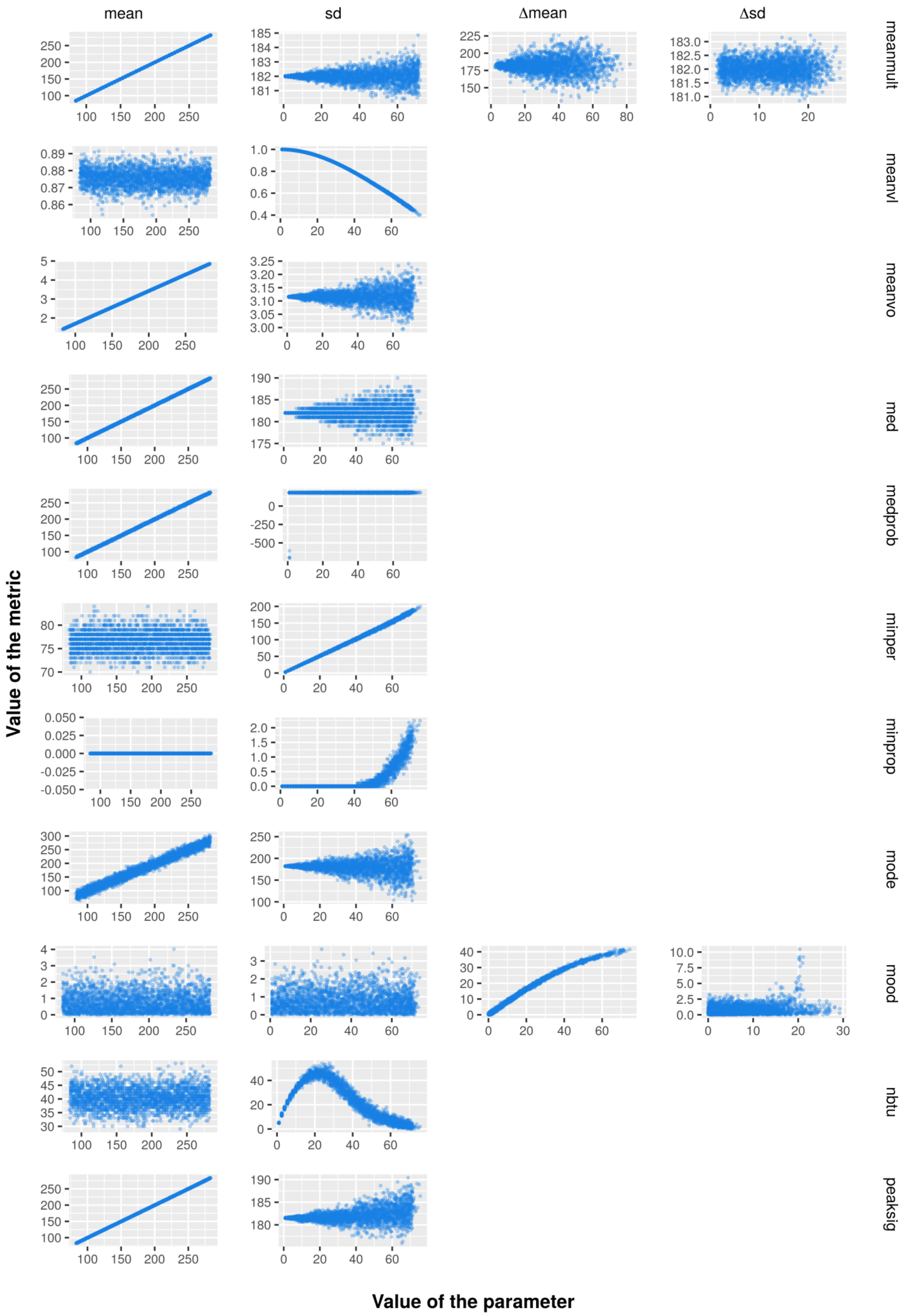
mean
sd
Δmean
Δsd
meanmult
meanvl
meanvo
med
medprob
minper
minprop
mode
mood
nbtu
peaksig
Value of the metric
Value of the parameter

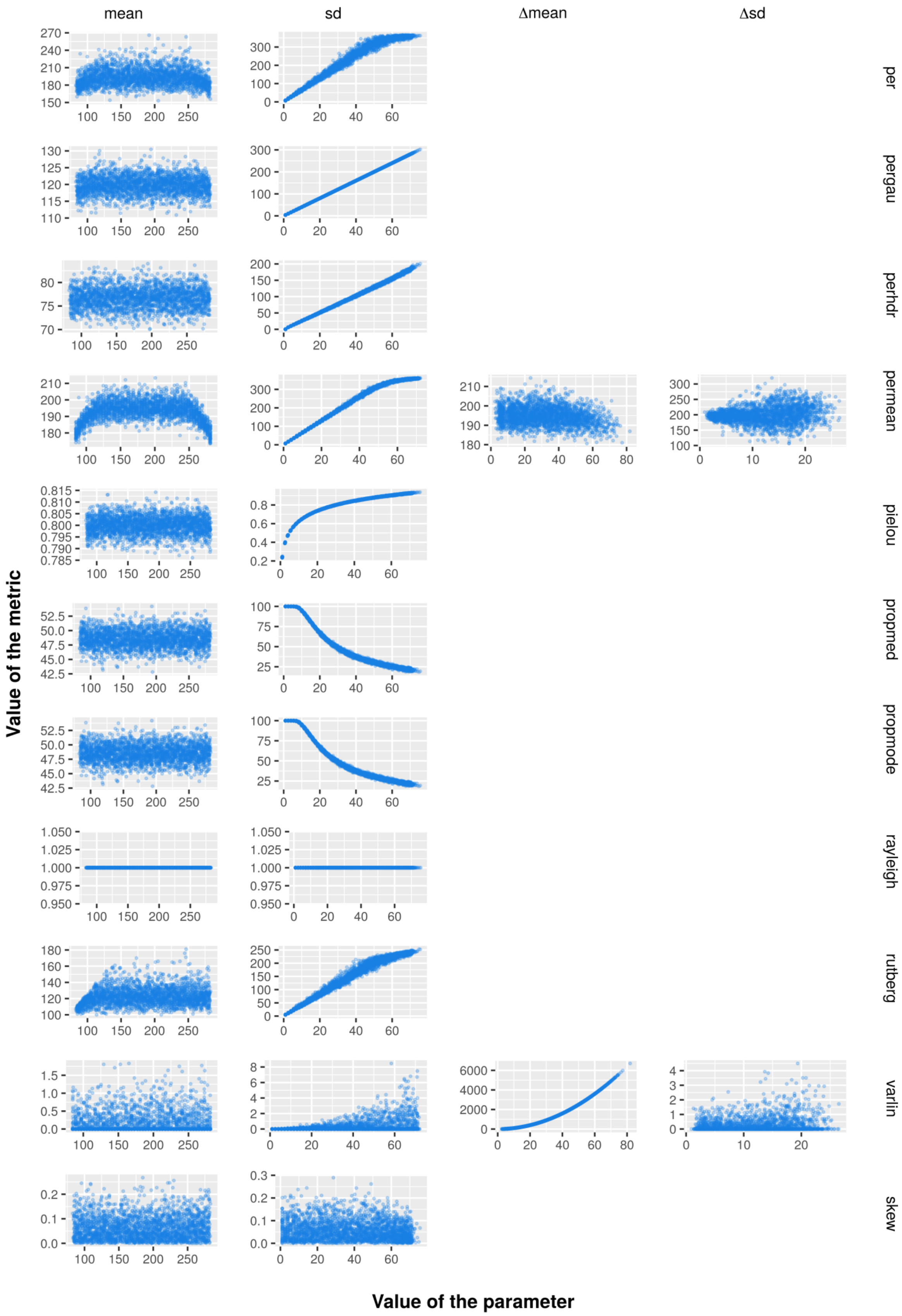
mean
sd
Δmean
Δsd
per
pergau
perhdr
permean
pielou
propmed
propmode
rayleigh
rutberg
varlin
skew
Value of the metric
Value of the parameter

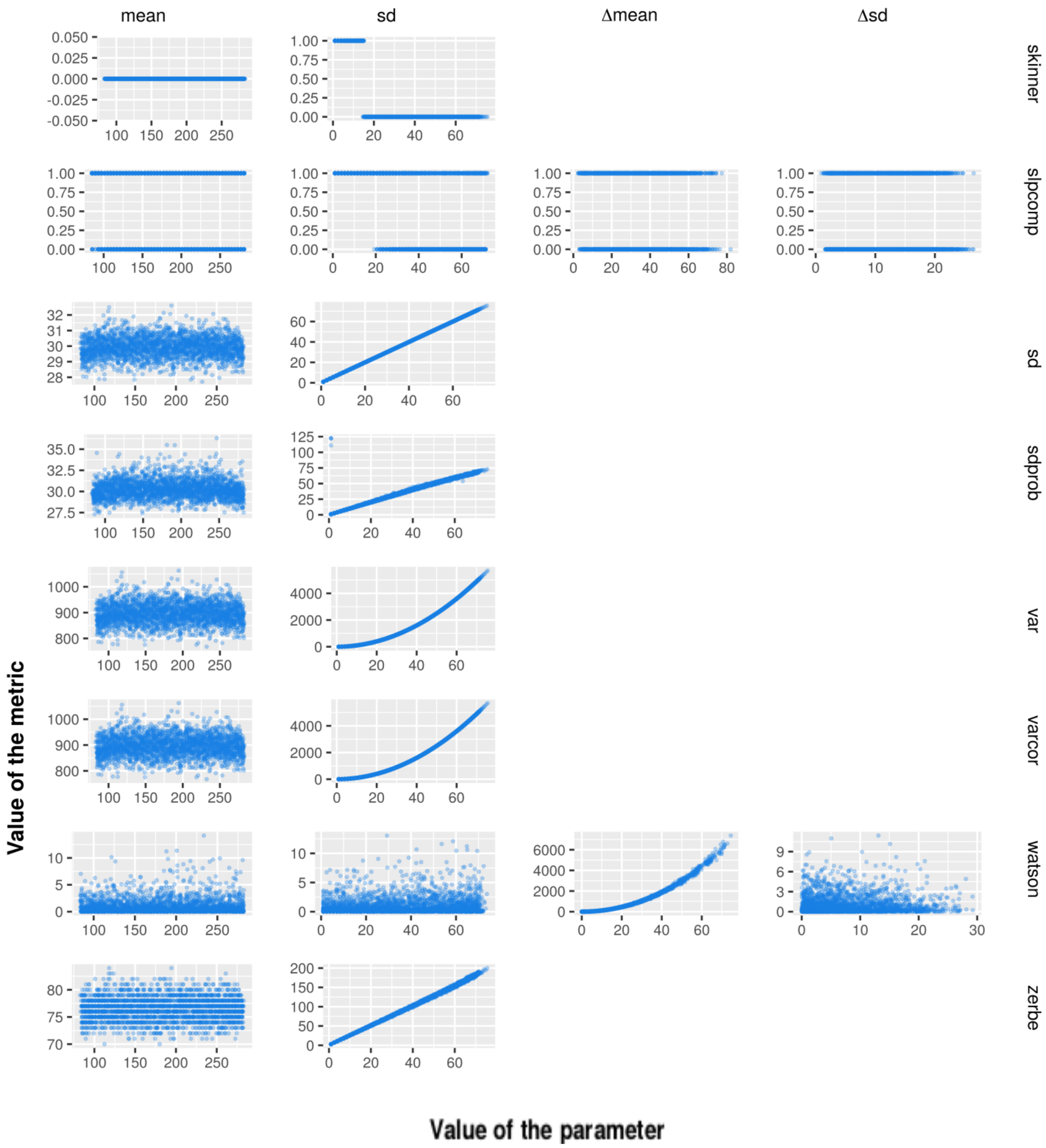
mean
sd
Δmean
Δsd
skinner
slpcomp
sd
sdprob
var
varcor
watson
zerbe
Value of the metric
Value of the parameter

Supporting information 6: Number of times each metric was used in the reviewed articles (*n* = 47). Inset: number of times each characteristic of phenology was studied in the reviewed articles, based on our *a priori* classification of each metric into one of the four characteristics we defined (see text for details: “Materials and methods” section, Step 1). *Green* = timing metrics, *orange* = synchrony metrics, *blue* = rhythmicity metrics, *pink* = regularity metrics.

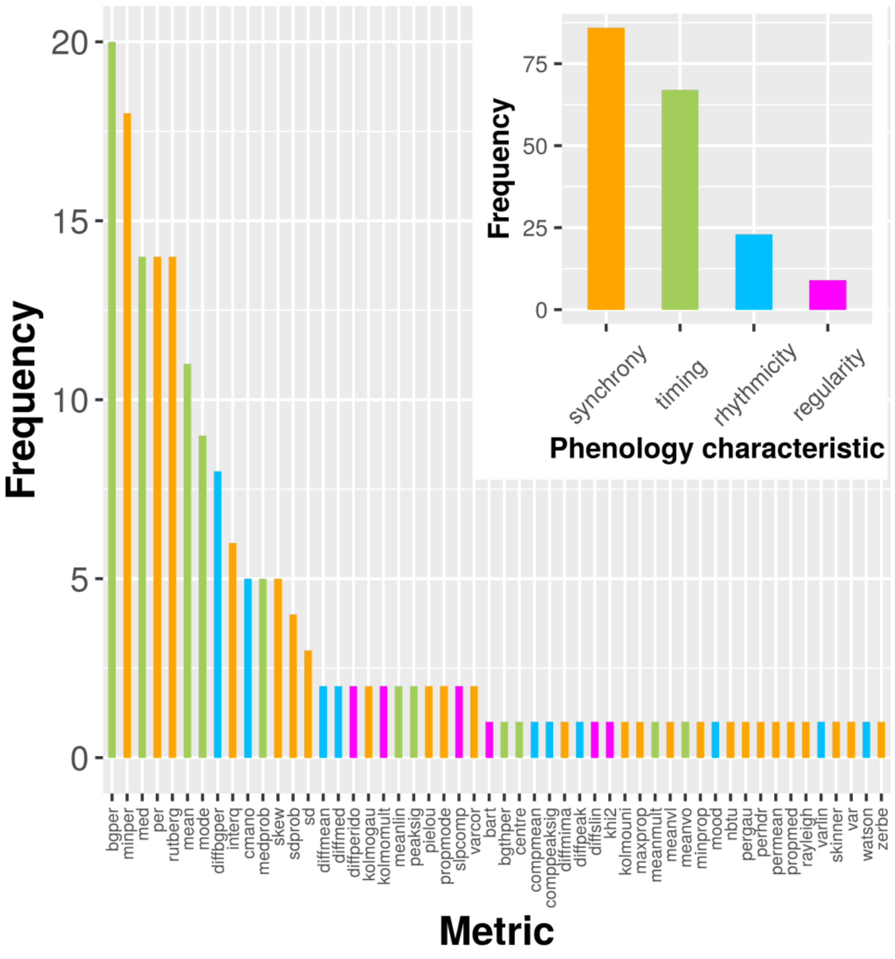

### References of the Supporting information:


Anderson, J. 1975. The occurrence of a secondary breeding peak in the southern impala. - Afr. J. Ecol. 13: 149-151.

Festa-Bianchet, M. 1988. Birthdate and survival in bighorn lambs (Ovis canadensis). - J. Zool. 214: 653-661.

Gaillard, J. M. *et al.* 1993. Timing and synchrony of births in roe deer. - J. Mammal. 74: 738–744.

Guinness, F. *et al.* 1978. Calving times of red deer (Cervus elaphus) on Rhum. - J. Zool. 185: 105-114.

Leuthold, W. and Leuthold, B. 1975. Temporal patterns of reproduction in ungulates of Tsavo East National Park. - E. Afr. Wildl. J. 13: 159-169.

Moe, S. R. *et al.* 2007. Trade-off between resource seasonality and predation risk explains reproductive chronology in impala. - J. Zool. 273: 237–243.

Rutberg, A. T. 1987. Adaptive hypotheses of birth synchrony in ruminants: an interspecific test. - Am. Nat. 130: 692–710.

Sinclair, A. R. E. *et al.* 2000. What determines phenology and synchrony of ungulate breeding in Serengeti? - Ecology 81: 2100–2111.
